# A self-generated Toddler gradient guides mesodermal cell migration

**DOI:** 10.1101/2021.12.16.472981

**Authors:** Jessica Stock, Tomas Kazmar, Friederike Schlumm, Edouard Hannezo, Andrea Pauli

## Abstract

The sculpting of germ layers during gastrulation relies on coordinated migration of progenitor cells, yet the cues controlling these long-range directed movements remain largely unknown. While directional migration often relies on a chemokine gradient generated from a localized source, we find that zebrafish ventrolateral mesoderm is guided by the uniformly expressed and secreted protein Toddler/ELABELA/Apela, acting as a self-generated gradient. We show that the Apelin receptor, which is specifically expressed in mesodermal cells, has a dual role during gastrulation, acting as a scavenger receptor to generate a Toddler gradient, and as a chemokine receptor to sense this guidance cue. Thus, we uncover a single receptor-based self-generated gradient as the enigmatic guidance cue that can robustly steer the directional migration of mesoderm through the complex and continuously changing environment of the gastrulating embryo.

**One sentence summary:** Aplnr has a dual role to self-generate and sense a Toddler gradient directing mesodermal cells during zebrafish gastrulation.

---

An animal’s body plan is first laid out during gastrulation which assembles the three germ layers—mesoderm, endoderm and ectoderm— through the combination of cell migration and differentiation^1–3^. While research throughout the last few decades has provided fundamental insights into how the germ layers are specified^3–6^, our knowledge of the molecular mechanisms that direct their spatial organization is very limited.

In zebrafish embryos, gastrulation starts with Nodal-induced specification and internalization of mesendodermal progenitor cells at the blastoderm margin^1,7,8^. Internalized cells subsequently migrate towards the animal pole, giving rise to the mesodermal and endodermal germ layers^3,9^. The molecular guidance, in particular of ventrolateral mesoderm migration, has remained largely unknown across vertebrates. The only factor currently known to be required for this process is the secreted protein Toddler/ELABELA/Apela^10,11^. Toddler is required primarily for mesoderm migration, acting through the Apelin receptor (in zebrafish, Apelin receptor a and b, collectively referred to as Aplnr), a G protein-coupled receptor specifically expressed in mesodermal cells^10–12^. However, how Toddler establishes directional migration during gastrulation has remained unclear.

Here, we show that mesodermal cells provide a dynamic sink to self-generate a Toddler gradient that acts as their elusive guidance cue. We find that Aplnr is the sole mediator of both Toddler gradient formation and sensing, which unveils a new mode of gradient self-generation in an *in vivo* system. Altogether, our study uncovers a simple yet robust mechanism that guides mesodermal cells through the dynamic and complex environment of a gastrulating embryo.

## RESULTS

### Toddler acts cell non-autonomously to attract Aplnr-expressing cells

We had previously shown that ventrolateral mesendoderm fails to migrate to the animal pole in *toddler* ^*-/-*^ embryos, yet the underlying cause remained unclear. To unveil the role of Toddler in mesendodermal cell migration, we first assessed the migratory behavior of mesendodermal progenitors. To this end, we transplanted between one and five marginal cells from a LifeAct-GFP-expressing wild-type or *toddler* ^*-/-*^ donor embryo to an unlabeled, stage- and genotype-matched host embryo and tracked cells after internalization using either light sheet (**Fig. 1A-C, fig. S1, movies S1-S3**) or confocal microscopy (**Fig. 1D-I, fig. S2, movies S4 and S5)**. Consistent with previous observations^2,10,13,14^, wild-type cells migrated in a directional manner from the margin toward the animal pole, were polarized, and extended actin-rich protrusions toward the animal pole (**Fig. 1A, D-H, fig. S1, movie S1**). In contrast, internalized *toddler* ^*-/-*^ cells in *toddler* ^*-/-*^ embryos displayed non-directional migration and were often dragged along with epiboly movements due to reduced and randomly oriented polarity and lamellipodia formation, concomitant with an increased frequency of cell blebbing (**Fig. 1B-I, fig. S1 and S2, movies S2 and S3**). Given that higher cell density can increase the frequency of blebbing^15–17^, this transition from actin- to bleb-based protrusions is likely a secondary effect that can be attributed to the increased cell density at the margin in *toddler* ^*-/-*^ mutant embryos. Taken together, these results reveal that, while *toddler* ^*-/-*^ cells are able to migrate in the absence of Toddler, they lack directionality towards the animal pole.

**Fig. 1.**
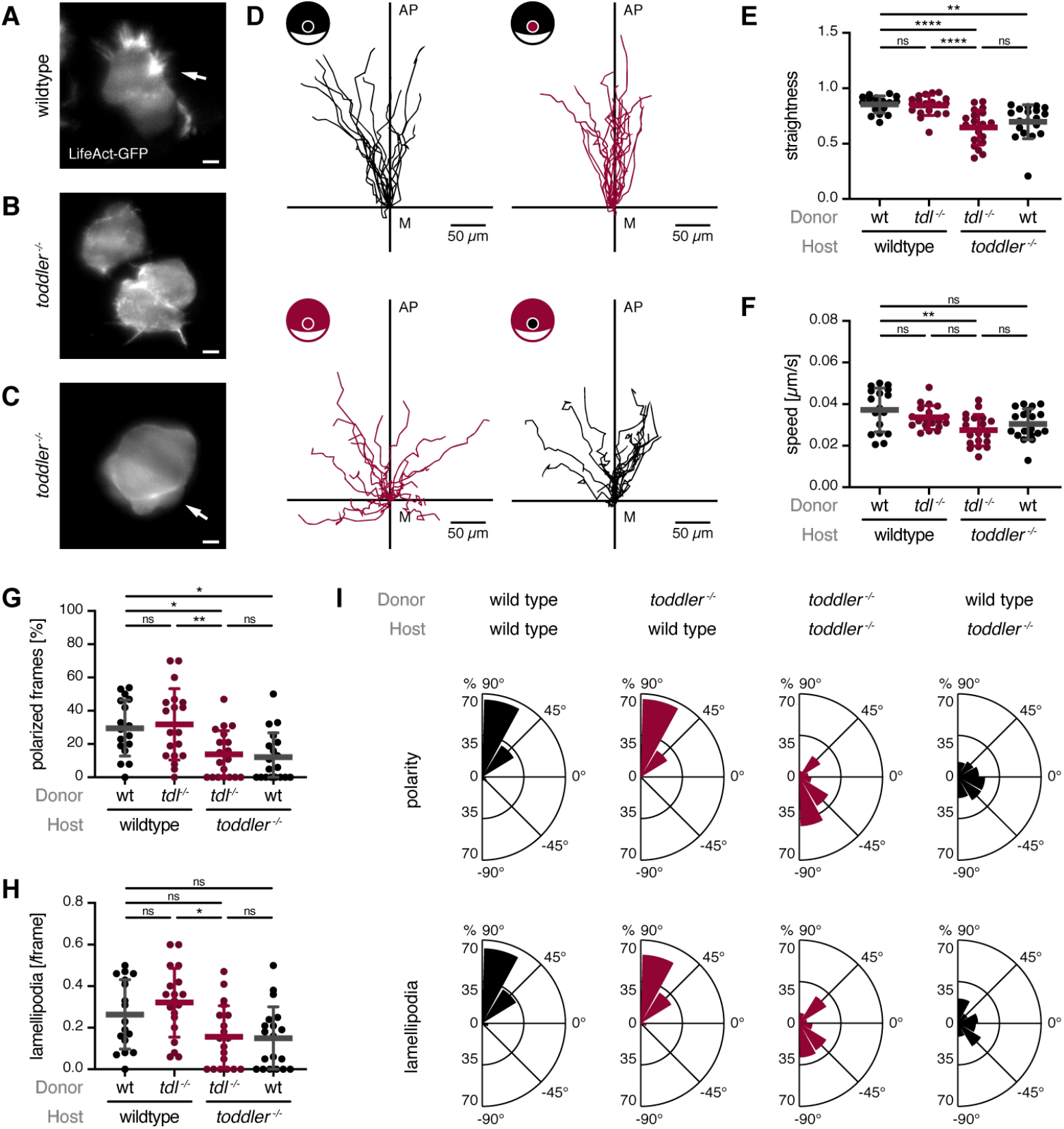
Toddler acts cell non-autonomously to mediate animal pole-directed polarization and migration of mesendodermal progenitors. Cell transplantation assays to assess the role and cell autonomy of Toddler signaling. LifeAct-GFP-labelled reporter cells were transplanted from the margin of a wild-type or *toddler* ^*-/-*^ donor embryo to the margin of a host embryo of the same or opposite genotype. **(A-C)** Light sheet microscopy images of representative internalized reporter cells. (**A**) Wild-type reporter cells in wild-type embryos extending actin-rich lamellipodia (white arrow) towards the animal pole. (**B-C**) *toddler* ^*-/-*^ reporter cells in *toddler* ^*-/-*^ embryos displaying lack of lamellipodia and polarization (B) or cell blebbing (C, white arrow). Scale bars, 10 μm. **(D)** Migration tracks of transplanted reporter cells (n = 19 cells for all conditions, except for wild type to wild type transplantation (n = 17)). Cells were tracked for 90 min after internalization using confocal microscopy. Genotypes of donor cells and host embryos are indicated in the embryo scheme (black: wild type; red: *toddler*^*-/-*^). x-axis = margin; y-axis = animal-vegetal axis; coordinate origin = start of track. **(E)** Quantification of track straightness calculated as the net displacement after 90 min divided by the total track length. **(F)** Quantification of migration speed of cells tracked in (D). **(G)** Quantification of cell polarity of internalized cells represented as the percentage of frames in which a cell was polarized (see Materials and Methods for classification of polarization). **(H)** Quantification of lamellipodia represented as average number of lamellipodia detected per frame (see Materials and Methods for classification of lamellipodia). **(I)** Rose plots showing relative enrichments (percentages) of orientations of polarity and lamellipodia, normalized to the total number of polarity axes or lamellipodia of all cells within the same condition. Data are means ± SD. Significance was determined using one-way ANOVA with multiple comparison; ****, p < 0.0001; **, p < 0.01; *, p < 0.05; n.s., not significant. Rose plots: 90° = animal pole; 0° = ventral/dorsal; −90° = vegetal pole. All graphs are oriented with the animal pole towards the top.

Given that Toddler is a secreted protein^10^, we hypothesized that it regulates directional migration of mesodermal progenitors via a cell non-autonomous mechanism. To distinguish cell intrinsic from extrinsic effects, we transplanted marginal cells from wild-type or *toddler* ^*-/-*^ donor embryos into unlabeled, stage-matched host embryos of the opposite genotype. Intriguingly, *toddler* ^*-/-*^ cells placed into a wild-type host embryo were able to migrate directionally, polarize and extend actin-rich protrusions towards the animal pole (**Fig. 1D-I, movie S6**). Wild-type cells in a Toddler-deficient environment, on the other hand, displayed the *toddler* ^*-/-*^ phenotype and failed to migrate directionally away from the margin (**Fig. 1D-I, fig. S2, movie S7**). Combined, these results confirm that Toddler acts cell non-autonomously to mediate the migration of mesendodermal progenitors.

A cell non-autonomous signaling mechanism, as well as the loss of directional migration and polarity in the absence of Toddler indicates that Toddler could act as a chemokine to guide mesodermal cells to the animal pole. As previous efforts to unravel the role of Toddler have led to contradicting results, providing support for a chemoattractant^18^ as well as a motogen^10^ function, we revistited this question and assessed the ability of Toddler to attract Aplnrb-expressing cells. Aplnrb-sfGFP-expressing cells from a *toddler* ^*-/-*^ donor embryo were transplanted next to a source of Toddler at the animal pole of a stage-matched *toddler* ^*-/-*^ host embryo (**Fig. 2A-B**). We found that Aplnrb-sfGFP-expressing cells displayed directional migration towards a Toddler-expressing source until they made initial contact with and attached to Toddler-expressing cells (**Fig. 2B-E, movie S8, see also Fig. S4M-N**). This cellular behavior is similar to the one described for the previously reported chemokine receptor/ligand pair Cxcr4b/Cxcl12a^19^. Directional migration was induced robustly over a distance of 100 μm (**Fig. 2C**) and required both the ligand and receptor to be present since the lack of either Toddler in the source or of Aplnr in the migrating cells (knockdown using *aplnra/b* morpholinos) led to the loss of directional migration (**Fig, 2B-E, movie S8**) and reduced contacts between Aplnrb-sfGFP-expressing and Toddler-expressing cells (**fig. S3, movie S8**). Taken together, these results indicate that Toddler can provide a chemoattractant signal and that Aplnr is necessary as a chemokine receptor to receive the signal.

**Fig. 2.**
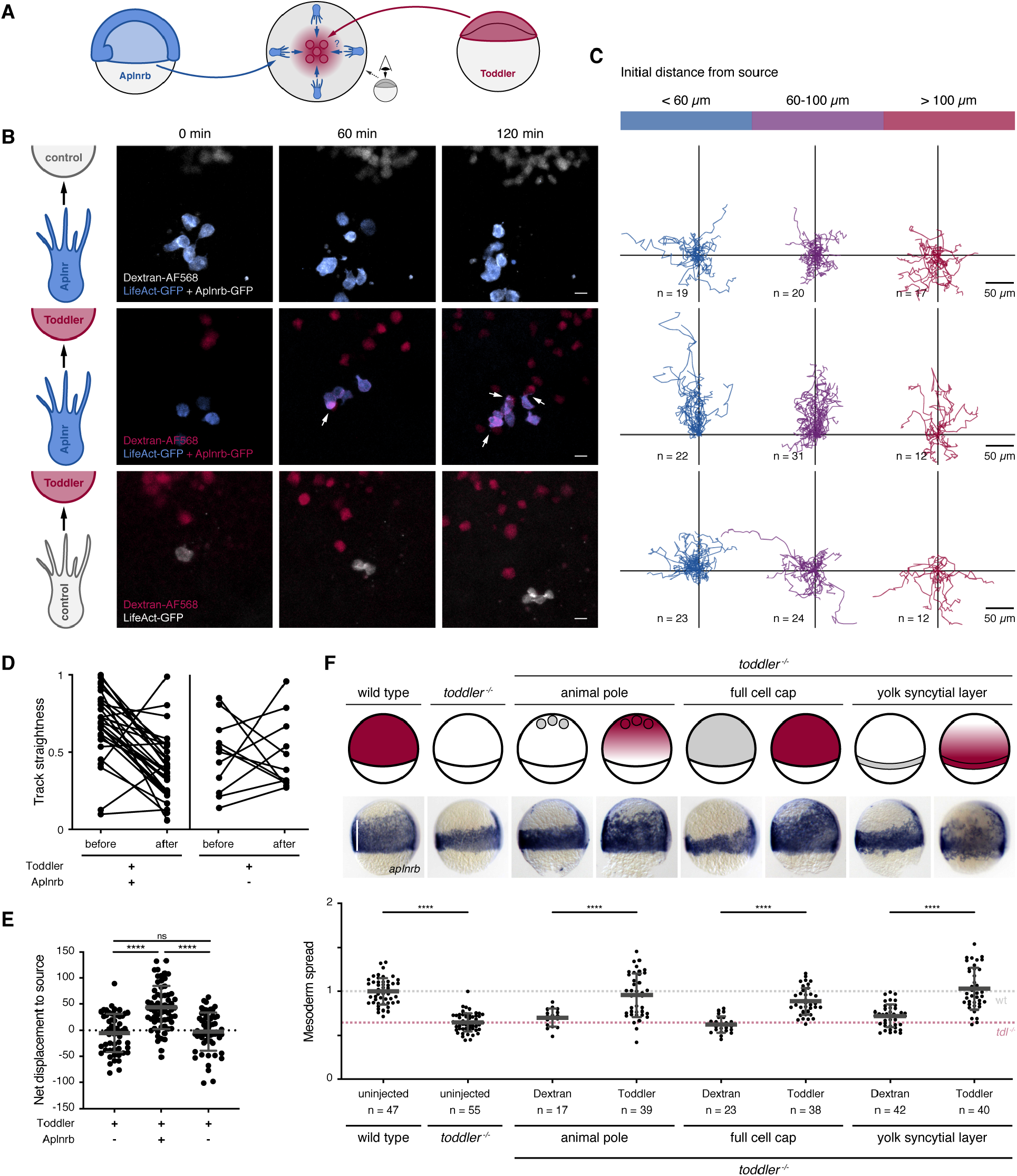
Aplnr-expressing cells are attracted towards a local source of Toddler. **(A)** Schematic representation of the experimental set-up to test for Toddler functioning as a chemokine signal for Aplnr-expressing cells. Toddler-expressing cells, red; Aplnrb-expressing cells, blue. **(B)** Snapshots of a time-laps confocal imaging series assessing ability of Aplnrb-sfGFP-expressing *toddler* ^*-/-*^ cells to react to an ectopically located Toddler or control source. (Top) Exposure of Aplnr-sfGFP-expressing cells (blue) to Toddler-deficient control cells (grey). n = 56 cells. (Middle) Exposure of Aplnr-sfGFP-expressing cells to Toddler-overexpressing source cells (red). n = 65 cells. White arrows indicate contact between Aplnrb-sfGFP-expressing cells and Toddler source cells. (Bottom) Exposure of cells deficient in Aplnr expression (grey) to Toddler-overexpressing source cells (red). n = 59 cells. **(C)** Cell tracks corresponding to conditions described in (B). Tracks were grouped by the distance of the cell from the source at the start of imaging (blue: >60 μm, purple: 60-100 μm, red: >100 μm). **(D)** Quantified track straightness of all Aplnrb-sfGFP-expressing and Aplnr-deficient cells that reach direct cell-cell contact with a Toddler-expressing source cell. Track straightness was compared before and after contact with the source cell. **(E)** Quantification of net displacement towards the source. **(F)** Ectopic expression of Toddler in *toddler* ^*-/-*^ embryos according to schematic representation of embryos at 75% epiboly (top). Animal pole source: transplantation of Toddler-overexpressing cells to the animal pole of sphere stage *toddler* ^*-/-*^ embryos; ubiquitous expression: injection of 2 pg *toddler* mRNA (rescuing concentration) into 1-cell stage *toddler* ^*-/-*^embryos; marginal source: injection of 10 pg of *toddler* mRNA into the yolk syncytial layer of 1k-cell stage *toddler* ^*-/-*^ embryos. Toddler (red); Dextran control (grey). Mesoderm spread was assessed using *in situ* hybridization for *aplnrb* (middle; lateral view, dorsal on the right; white line indicates measurement of ventral mesoderm spread) and quantified relative to the average spread in wild-type embryos (bottom). Data are mean ± SD. Significance was determined using one-way ANOVA with multiple comparison; ****, p < 0.0001; **, p < 0.01; n.s., not significant. Scale bars, 20 μm. All graphs and images are oriented with the source at the top.

### Mesodermal cells are guided by a self-generated Toddler gradient

Our results so far indicate that Toddler can act as a guidance cue that attracts mesodermal cells. In support of this, placing a cluster of Toddler-expressing cells at the animal pole is sufficient to restore animal pole-directed mesoderm migration in *toddler* ^*-/-*^ embryos (**Fig. 2F**). However, in line with previous results^10^, expression of Toddler throughout the embryo cap, leading to its homogenous distribution, or from the margin (*toddler* mRNA injection into the yolk syncytial layer (YSL) before the onset of gastrulation), causing Toddler to be graded in the opposite direction, are also able to rescue animal pole-directed mesoderm migration in *toddler* ^*-/-*^ embryos (**Fig. 2F**). These results are contrary to typical chemokine models based on concentration gradients formed by a fixed source or sink, and suggest that mesoderm migration during gastrulation requires Toddler, but is largely independent of Toddler’s expression site.

Studies in the zebrafish lateral line^20,21^ and *Dictyostelium*^22,23^ have shown that migrating cells can “self-generate” chemokine gradients by locally taking up the chemokine, making them independent of a localized source. To assess quantitatively whether mesodermal cells could self-polarize by locally shaping a Toddler gradient, we turned to computational modeling (see **Materials & Methods**). We simulated in time the one-dimensional migration of mesodermal cells along the animal-vegetal axis *x* with local cell density *c(x,t)* (**Fig. 3A**, see also **fig. S4A-E** for a sensitivity analysis and **Materials & Methods** for details). Cells internalize at the margin (*x=0*), can randomly diffuse (random movements with diffusion coefficient *D*_*c*_), and migrate directionally according to the local gradient of the Toddler concentration profile *T(x,t)*. Simulation of the Toddler concentration *T(x,t)* along the animal-vegetal axis factored in random Toddler diffusion (with coefficient *D*_*T*_), baseline degradation (with timescale *τ*_*T*_), and a sink to remove Toddler that is proportional to the local density of mesodermal cells (see **Materials & Methods** for details). Importantly, we also took into consideration that the quantity of available Toddler is not a fixed value, as in typical *in vitro* assays of self-generated gradients^22,23^. Indeed, based on single-cell RNA-seq data^24^, *toddler* mRNA is expressed ubiquitously throughout the embryo at the onset of gastrulation (**Fig. 3B**) and becomes restricted to non-mesodermal cells during gastrulation (indicated by the mutually exclusive expression patterns of *toddler* and *aplnrb* (**Fig. 3B-C**)). Based on these observations, we infer that Toddler is continuously and homogenously expressed along the animal-margin axis (e.g., by the overlying ectodermal tissue) throughout gastrulation.

**Fig. 3.**
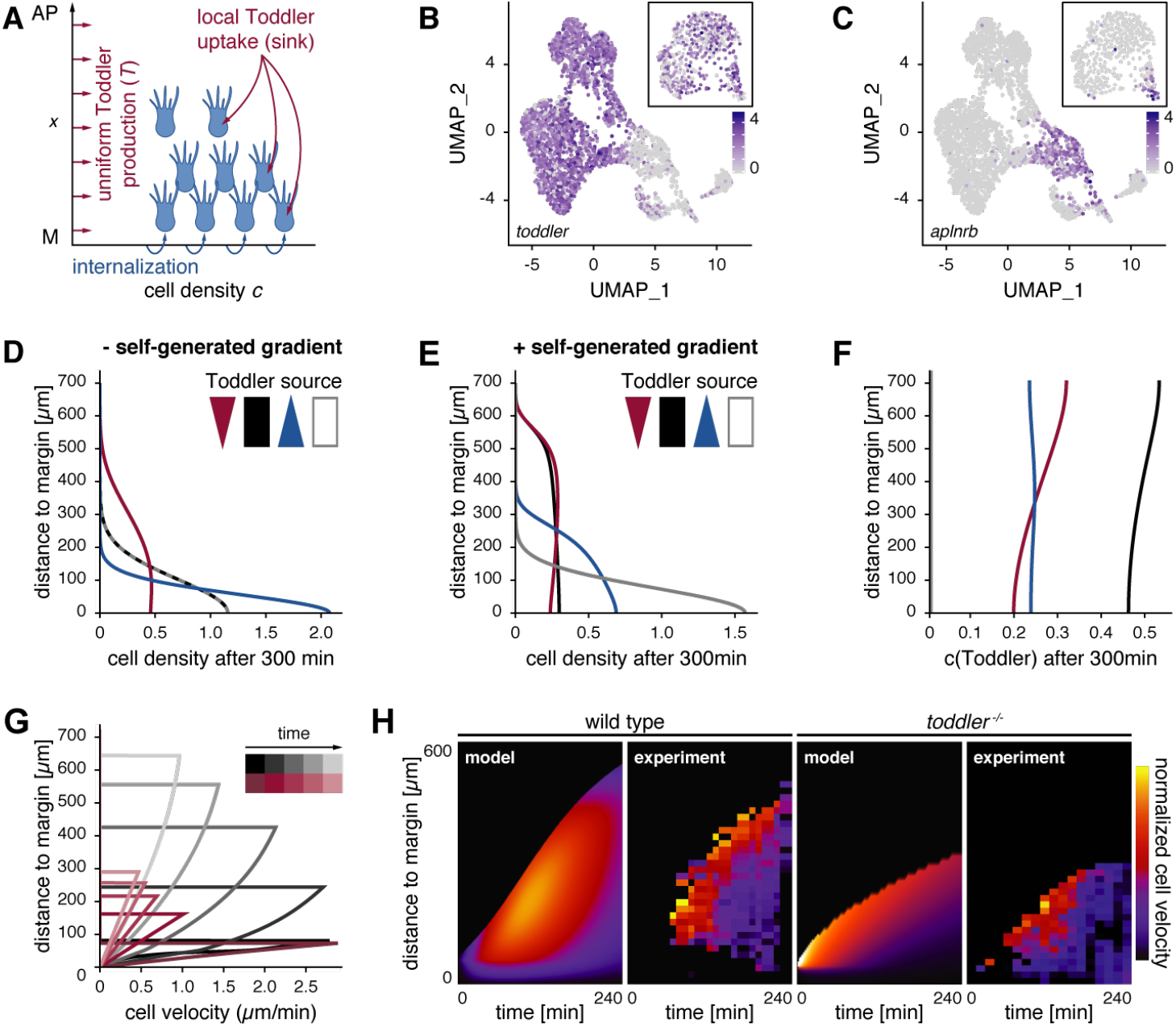
Computational simulations predict a self-generated Toddler gradient. **(A)** Schematic representation of the one-dimensional model of mesoderm density and Toddler concentration along the animal-vegetal axis (x = 0 denotes the margin, at which mesodermal cells are added). Toddler (red) is produced uniformly at rate T_0_ and is degraded locally by mesoderm cells (blue). Mesodermal cells can move randomly as well as directionally in response to local Toddler gradients. **(B-C)** Uniform manifold approximation and projection (UMAP) clustering of single cells at 60% epiboly based on single cell RNAseq data ^24^. Inset depicts UMAP at 30% epiboly. Color-code represents expression levels of *toddler* (B) and *aplnrb* (C) in individual cells, respectively. (**D-E**) Predicted mesoderm density profiles (arbitrary units) after 300 min without (D) or with (E) Toddler uptake by mesoderm cells, for different profiles of Toddler production T_0_(x): graded towards the margin (blue), graded towards the animal pole (red), uniform (black) or no production (white/grey). **(F)** Predicted Toddler concentrations (arbitrary units) after 300 min with Toddler uptake by mesodermal cells. Profiles of Toddler productions T_0_(x) as described in (D and E). **(G)** Predicted spatiotemporal profiles of mesodermal cell velocities in wild-type (black) and *toddler* ^*-/-*^ embryos (red). **(H)** Predicted (model) and experimental (experiment) kymographs of mesodermal cell migration in wild-type (left) and *toddler* ^*-/-*^ (right) embryos. Experimental data from light-sheet microscopy and tracking of *drl:GFP*-positive cells (N = 7 (wild type) and N = 6 (*toddler* ^*-/-*^) embryos; average number of n = 195 cells tracked per embryo). Color-code represents normalized velocity (yellow: high; dark purple: low).

To test the core assumptions of the model and constrain its parameters, we first measured the baseline degradation time *τ*_*T*_ of Toddler. To this end, we injected *in vitro* synthesized Toddler peptide into *toddler* ^*-/-*^, *MZoep* ^*-/-*^ double mutant embryos deficient for both Toddler production (*toddler* ^*-/-*^) and the proposed mesodermal Toddler sink (*MZoep* ^*-/-*^)^25,26^. We found that a single time scale can describe Toddler degradation kinetics in *MZoep* ^*-/-*^, *toddler* ^*-/-*^ double mutants, from which we can extract the parameter *τ*_*T*_ ≈ 2h (**fig. S4F-G**). Given the known molecular weight of Toddler, we could also estimate its random diffusion to be *D*_*T*_ ≈ 3.10^3^*μm*^2^/*min* (a typical value for small diffusible molecules, see Müller et al., 2012 and **Materials & Methods**). Based on these calculations, locally produced Toddler can propagate up to length scales of 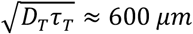 (the typical size of a zebrafish embryo) in the absence of a mesoderm sink. Altogether, these measurements provided estimates for the key parameters of Toddler dynamics, which we used to make predictions on spatiotemporal dynamics of mesoderm migration.

We simulated wild-type migration in the presence or absence of the mesodermal Toddler sink. The simulation predicted that animal pole-directed migration of mesodermal cells would be abolished in the absence of the sink function, if the Toddler gradient were absent (ubiquitous expression or *toddler* ^*-/-*^) or reversed (preferential expression at the margin from the YSL) (**Fig. 3D**). With sink function in mesodermal cells, the dynamics depended on one rescaled parameter which represents the strength of the sink function and the strength of the polarization response to a given Toddler gradient (see **Materials & Methods**). Introduction of this parameter resulted in local depletion of Toddler at the margin, where mesodermal cells internalize, thus forming a Toddler concentration gradient (**Fig. 3E-F**). The resulting animal pole-directed migration of mesodermal cells sustains the Toddler gradient, with stronger sink function resulting in proportionally faster migration and higher independence of the location of the Toddler source. Thus, in agreement with our experimental findings, the self-generated gradient model suggests that mesodermal cells can shape a gradient from uniform Toddler concentrations (or even from an inverse Toddler gradient, as long as the sink function is strong enough) to direct their migration to the animal pole (**Fig. 3E**). Importantly, the resulting Toddler gradient can be sustained even close to the margin (far from the migratory leading edge) due to the continuous production of Toddler.

This model further predicts a distinct pattern of velocities across mesodermal cells, in which cell velocities increase as a function of the distance from the margin but decrease overall as a function of time (**Fig. 3G**). To test this prediction experimentally, we used light sheet microscopy to image transgenic zebrafish embryos expressing *drl:GFP* (*draculin* promoter driving GFP expression), which specifically labels ventrolateral mesoderm during gastrulation^12,28^, injected with *h2b-RFP* mRNA to label all nuclei. This reporter allowed us to identify and track ventrolateral mesodermal cells and their progenitors (see **Materials and Methods** for details). We found that the experimentally measured average mesodermal cell velocity was indeed highest at the leading edge of the mesodermal cells and decreased overall with the progression of gastrulation (**Fig. 3H**). Therefore, with a single fitting parameter (strength of self-generated directionality as mentioned above, see **Materials & Methods** for details), theoretical predictions could recapitulate the experimental kymographs (**Fig. 3H**). Moreover, the shape of the mesodermal cell density profile along the animal-vegetal axis in wild-type embryos predicted by the model (**fig. S4H**) was also consistent with experimental data (**fig. S4I-J**).

Next, we simulated mesodermal migration dynamics in the absence of Toddler. In this case, the only mode of mesodermal cell movements is random cell motility (with coefficient *D*_*c*_ estimated from short-term measurements of transplanted cell displacements in *toddler* ^-/-^ embryos, see **fig. S4K-L**). The model predicted that mesodermal cells migrate approximately only half the distance in *toddler* ^*-/-*^ embryos compared to that in wild-type embryos, and that animal pole-directed migration rapidly decreased over time (**Fig. 3G-H**), as expected from a random migration process. Indeed, we found that the migration front of mesodermal cells in *toddler* ^*-/-*^ *tg(drl:GFP)* embryos did not progress as far as in wild-type *tg(drl:GFP)* embryos and had nearly zero net velocity four hours after the onset of internalization (**Fig. 3H**).

In summary, our theoretical predictions, supported by experimental measurements in wild-type and *toddler* ^*-/-*^ embryos, reveal that mesodermal cells are guided to the animal pole by a self-generated Toddler gradient.

### Aplnr-mediated removal of Toddler at the margin self-generates a Toddler gradient

In our model, mesodermal cells are required to remove Toddler locally in order to self-generate a Toddler concentration gradient. To test this hypothesis, we transplanted LifeAct-GFP-labelled cells from the margin of a wild-type embryo to the margin of stage-matched host embryos that either possess (wild-type) or lack (*MZoep* ^*-/-*^) the proposed mesodermal sink (**Fig. 4A**). The reporter cells showed directional migration in the wild-type host but lost their directionality and polarity in the *MZoep* ^*-/-*^ host (**Fig. 4B-F, fig. S5**). Instead, reporter cells in the absence of a mesoderm sink formed multiple actin-rich protrusions in various directions (**Fig. 4C and E-F, fig. S5, movie S9**), in line with excessive chemokine stimulation from all directions^29^.

**Fig. 4.**
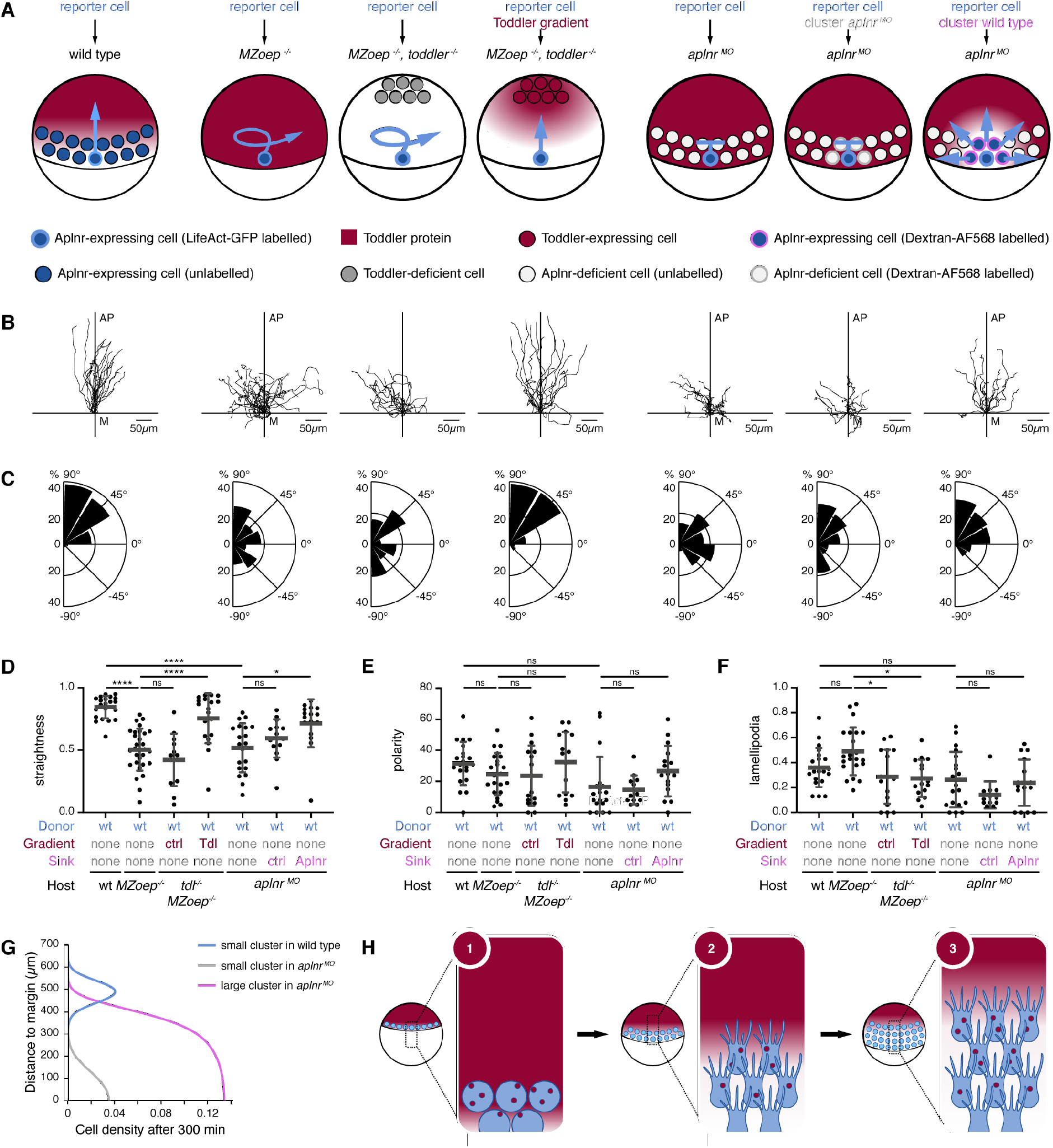
Aplnr-expressing mesodermal cells are required to establish a Toddler gradient. Cell transplantation assays to test for the necessity of Aplnr-expressing mesodermal cells as a sink for Toddler in the gastrulating embryo. Transplanted LifeAct-GFP-labelled wild-type reporter cells were used as a read-out for the presence and shape of a Toddler gradient. **(A)** Schematic representation of the different scenarios tested. From left to right: (1) Transplantation of reporter cells into a wild-type host embryo (n = 21). (2) Transplantation of reporter cells into an *MZoep* ^*-/-*^ host embryo (n = 22). (3) Transplantation of reporter cells into an *MZoep* ^*-/-*^, *toddler* ^*-/-*^ double mutant host embryo. Additional transplantation of Dextran-AlexaFluor568-labelled control source cells to the animal pole (n = 16). (4) Transplantation of reporter cells into an *MZoep* ^*-/-*^, *toddler* ^*-/-*^ double mutant host embryo. Additional transplantation of Toddler-expressing source cells to the animal pole to simulate presence of a Toddler gradient (n = 15). (5) Transplantation of reporter cells into an *aplnr* ^*MO*^ embryo (n = 20). (6) Co-transplantation of 1 to 5 reporter cells and a large number of Dextran-AlexaFluor568-labelled Aplnr-deficient control cells into *aplnr* ^*MO*^ host embryo (n = 15). (7) Co-transplantation of 1 to 5 reporter cells and a large number of Aplnr-expressing cells into *aplnr* ^*MO*^ host embryos to re-introduce a localized Toddler sink (n = 16). Reporter cell is depicted in dark blue with light blue rim. Expected Toddler gradient is represented in red. Blue arrow indicates expected migration behavior of the reporter cells. **(B)** Tracks of transplanted reporter cells (order as described in (A)). Cells were tracked for 90 min after internalization. x-axis = margin; y-axis = animal-vegetal axis; coordinate origin = start of track. **(C)** Rose plots showing relative enrichments (percentages) of orientations of polarity, normalized to the total number of polarity axes of all cells within the same condition. 90° = animal pole; 0° = ventral/dorsal; − 90° = vegetal pole. **(D)** Quantification of straightness of cell tracks presented in (B). **(E)** Quantification of cell polarity of reporter cells represented as percentage of frames in which a cell was polarized. **(F)** Quantification of lamellipodia detected in reporter cells per frame. **(G)** Numerical simulation of scenarios presented in (A) with the same model parameters as in Figure 3. Migration of small or large clusters of Aplnr-expressing mesodermal reporter cells, which are able to take up Toddler, was simulated in the presence (wild type) or absence (*aplnr*^MO^) of Aplnr in host embryos (all other parameters being identical). **(H)** Schematic representation of the self-generated Toddler gradient: (1) Toddler is ubiquitously expressed throughout the embryo cap during zebrafish gastrulation. Mesodermal progenitor cells internalize at the margin and express Aplnr, which acts as a scavenger receptor for Toddler. (2) Aplnr binds and internalizes Toddler, which generates a local Toddler concentration gradient in front of the mesodermal cells, providing a directional cue. (3) Aplnr also acts as a chemokine receptor to sense the self-generated Toddler guidance cue and induces the directed migration of mesodermal cells towards the animal pole, while continuously internalizing Toddler and shaping the local gradient. Data are means ± SD. Significance was determined using one-way ANOVA with multiple comparison; ****, p < 0.0001; *, p < 0.05; n.s., not significant. All images and graphs are oriented with the animal pole towards the top.

To confirm that the lack of a Toddler gradient in the absence of mesendodermal cells causes the loss of directional migration, we engineered a Toddler gradient in *MZoep* ^*-/-*^, *toddler* ^*-/-*^ double mutant embryos. For this purpose, we transplanted a cluster of Toddler-expressing cells to the animal pole of these embryos to establish a local source of Toddler (**Fig. 4A**). Strikingly, this ectopically induced Toddler gradient was able to rescue directional polarization, protrusion formation, and migration of wild-type reporter cells towards the animal pole in embryos lacking mesendodermal cells (**Fig. 4B-F, movie S9**). Therefore, the mesendodermal cell population in the embryo serves as a sink that removes Toddler locally at the margin, generating a gradient of increasing Toddler concentration towards the animal pole which ultimately drives directional mesodermal cell migration.

Although only a few examples of self-generated gradients have been described, different mechanisms have been observed to mediate their formation, such as enzymatic chemokine digestion at the cell surface^22^ or chemokine internalization via a scavenger receptor^20,21^. Toddler binds Aplnr-expressing cells^11^ and induces Aplnr internalization^10^ (**fig. S6**), which, based on the general mechanism of GPCR-ligand interaction and signaling^30^, should lead to the concomitant endocytosis and degradation of Toddler. To test whether Aplnr acts as a scavenger receptor for Toddler, we compared the stability of the injected Toddler peptide in *MZoep* ^*-/-*^, *toddler* ^*-/-*^ double mutant embryos to *toddler* ^*-/-*^ embryos either with normal mesoderm specification (normal sink) or upon overexpression of Aplnrb (enhanced sink). The Toddler degradation rate in embryos with endogenously expressed Aplnrb (*toddler* ^*-/-*^ embryos) was only very slightly increased in comparison to *MZoep*^*-/-*^, *toddler* ^*-/-*^ double mutants, while overexpressing Aplnrb in the entire embryo led to an increase in degradation rate. This is consistent with the locally restricted expression of endogenous Aplnrb in the small number of mesodermal cells at the margin (**fig. S4F-G**, see **Materials & Methods** for details). Thus, Aplnr is an excellent candidate to mediate mesoderm-dependent degradation of Toddler and function as a sink to self-generate a Toddler gradient.

To test our hypothesis that Aplnr in mesodermal cells acts as the scavenger receptor for Toddler, we transplanted individual LifeAct-GFP-labelled wild-type reporter cells to the margin of a sphere-stage *aplnr* ^*MO*^ host embryo, which lacks Aplnr expression in mesodermal cells (**Fig. 4A**). The reporter cells lost their ability to polarize, lacked the typical animal pole-directed bias in protrusion formation, and failed to migrate towards the animal pole in the *aplnr* ^*MO*^ host, phenocopying their behavior in the *MZoep* ^*-/-*^ host (**Fig. 4B-F, movie S10**). Based on these results, we propose a dual role for Aplnr in mesodermal cells during zebrafish gastrulation: (1) as a scavenger receptor to generate the Toddler gradient by removing it from the environment, and (2) as a chemokine receptor to sense the Toddler gradient and direct cell migration.

In light of the dual role of Aplnr, we expanded the computational model described above to account for the Aplnr turnover during Toddler removal. Due to the finite capacity of Aplnr to remove Toddler, the model generally predicts that, above a threshold level of Toddler, mesodermal migration would be non-directional, resembling *toddler* ^*-/-*^ embryos (**fig. S7A**). It further predicts that these migration defects at high Toddler levels can be rescued by simultaneous overexpression of Aplnr (**fig. S7B**). Importantly, we confirmed both of these predictions experimentally. Firstly, overexpression of Toddler in wild-type embryos caused mesodermal mis-migration phenotypes mimicking *toddler* ^*-/-*^ embryos (**fig. S7C**), as previously observed (Pauli et al., 2014). Secondly, simultaneously increasing the number of mesodermal cells (and thereby the level of Aplnr expression in these embryos) by blocking the expression of the Nodal inhibitor Lefty2^31^ restored mesoderm migration in embryos with intermediate levels of Toddler overexpression (5 pg *toddler* mRNA, **fig. S7C**). Taken together, these observations provide independent evidence for Aplnr acting as a scavenger receptor for Toddler and support the model of a self-generated gradient.

Finally, the formation of a self-generated gradient requires a sufficient number of cells to remove enough chemokine locally to establish the gradient^32^, as seen by a decrease of animal pole-directed migration in simulations with a reduced number of mesodermal cells (**Fig. 4G**). Therefore, we hypothesized that a cluster of Aplnr-expressing cells, but not a single Aplnr-expressing cell, would undergo directional migration in a uniform Toddler environment when transplanted to the margin. To test this hypothesis, we assessed the migratory ability of a small number (up to 10) of LifeAct-GFP-labelled wild-type reporter cells co-transplanted with a large number (more than 50) of Dextran-Alexa568-labelled, Aplnr-expressing (sink^+^) or Aplnr-lacking (sink^-^) cells, to the margin of an *aplnr* ^*MO*^ host embryo (**Fig. 4A**). The reporter cells migrated randomly and failed to reach the animal pole when co-transplanted with cells that lacked Aplnr (**Fig. 4B-F, movie S10**). However, when the co-transplanted cells expressed Aplnr, the reporter cells displayed increased track straightness, reduced ectopic protrusion formation and increased directional, radial outward-directed polarization (**Fig. 4B-E, movie S10**), indicating the presence of a guidance cue. In summary, these results provide evidence that collectively a cluster of Aplnr-expressing cells is sufficient to restore sink activity and directional migration of reporter cells, in line with collective cell migration being a hallmark of self-generated gradients.

## DISCUSSION

This study describes a previously unknown, self-generated Toddler gradient, that is formed and sensed by a single molecular mediator, the Apelin receptor, to guide the animal pole-directed migration of ventrolateral mesoderm during zebrafish gastrulation (**Fig. 4H**). Through the global production and local degradation of Toddler, the location of the mesoderm sink defines the directionality of cell migration toward the animal pole, as it both generates and senses a Toddler gradient that is self-sustained and inversely proportional to the mesodermal cell density (**Fig. 4H**).

Different types of self-generated gradients have recently been discovered as powerful guidance mechanisms^20–22,33^. Self-generated chemokine gradients are characterized by one shared feature: Both the function of chemokine gradient formation and sensing are found within the migrating unit at opposite poles of the tissue^20,21^ or even the same cell, as suggested by theoretical work and studies with cultured cells *in vitro*^22^. Additional studies based on computational modeling^34^ or *in vitro* experiments^35^ have further suggested that removal of the chemokine and sensing of the gradient could be mediated by the same receptor. We provide experimental *in vivo* evidence that Aplnr indeed executes both scavenger and sensor roles, acting as the sole molecular player in generating and sensing the directional cue that guides mesodermal cells to the animal pole.

The use of a self-generated gradient, in particular those based on a single receptor, holds several advantages over a pre-existing gradient for the migration of a dynamic tissue, like the arising mesoderm. First, self-generated gradients can robustly act over long distances^32^, which is necessary for mesodermal cell migration, with the space from margin to animal pole spanning over 500 μm during zebrafish gastrulation. Secondly, the simple two-component system (here: Aplnr and Toddler) of a single receptor-based self-generated gradient can be easily adjusted to a dynamically changing tissue, and thus circumvents the need to establish stable domains with distinct functions. Instead, the shape of the gradient is determined by the position of the migrating cell front rather than the total size of the tissue. This is particularly beneficial during gastrulation, as the continuous internalization of mesodermal cells and the vegetal pole-directed movement of the margin causes a steady increase in cell number and expansion of tissue size. Finally, while pre-existing gradients require tight regulation of steepness and chemokine concentration along the gradient to ensure reliable cell guidance, self-generated gradients can compensate for changes in length scale (as described above) and chemokine levels by adjusting the rate of chemokine breakdown ^36^. Therefore, self-generated gradients may have evolved to accommodate different architectures of complex migrating tissues by providing the necessary flexibility to adjust to changing environments and migration modes.

In the future it will be interesting to establish tools to directly visualize the Toddler gradient in the embryo. This task has so far proven as technically challenging. Tagging of Toddler renders the protein non-functional and there is currently no antibody available that can detect Toddler at endogenous levels in immunostainings. Furthermore, the downstream signaling pathways in the embryo remain unclear, precluding the use of a reporter-based read-out. Nonetheless, our computational simulations and experimental analyses of mesodermal cell migration patterns are consistent with the formation of a Toddler gradient in wild-type embryos since in the absence of Toddler as a guidance cue mesodermal cell migration was mostly driven by random cell motility. However, it is important to note that *toddler* ^*-/-*^ cells retained some residual bias for migration towards the animal pole (**Fig. 1 and 3**). These findings suggest that—at least in the absence of a Toddler–based guidance cue—migration is aided by additional, Toddler-independent mechanisms, which could include Contact inhibition of locomotion^37,38^ or biomechanical forces^39,40^.

Morphogenetic movements during gastrulation across the animal kingdom share common principles. In amniotes, including mouse and human, mesodermal progenitors internalize at the primitive streak before migrating anteriorly as a non-coherent cell sheet ^1^. Previous studies have shown that, while conserved, Toddler and Apelin, the other ligand of Aplnr, are not essential for gastrulation movements in mice ^41^. However, based on the presented advantages and similar nature of morphogenetic movements across species, we hypothesize that self-generated gradients of different or redundant chemokine receptor-ligand pairs could present a universal mechanism underlying the anterior migration of mesodermal progenitor cells in gastrulation, as well as other cell migration events throughout development and physiology.

## ACKNOWLEDGMENTS

We thank Karin Aumayer and the team of the biooptics facility at the Vienna Biocenter, in particular Pawel Pasierbek and Tobias Müller, for support with microscopy; Karin Panser, Carina Pribitzer and the animal facility personnel for taking care of zebrafish; Mirjam Binner and Anna Bandura for help with genotyping; Tiago Lubiana Alves for sharing the code for scRNA-Seq analyses; the Heisenberg lab, in particular Diana Pinheiro, for joint lab meetings, discussions on the project and providing the *tg(gsc:CAAX-GFP)* fish line; the Raz lab for providing the Lifeact-GFP plasmid; Angela Andersen, Alex Schier, Carl-Phillip Heisenberg and Elly Tanaka for comments on the manuscript; the entire Pauli lab, in particular Krista Gert and Victoria Deneke, for valuable discussions and feedback on the manuscript.

## AUTHOR CONTRIBUTION

Conceptualization: JS, EH, AP

Methodology: JS

Data analysis: JS

Cell tracking analysis: TK, FS

Computational Modeling: EH

Funding acquisition: AP, EH

Supervision: AP

Writing – original draft: JS

Writing – review & editing: JS, AP, EH

## RESOURCE AVAILABILITY

Light sheet imaging data are available upon request.

## MATERIALS AND METHODS

### Ethical statement

All fish experiments were conducted according to Austrian and European guidelines for animal research and approved by the Amt der Wiener Landesregierung, Magistratsabteilung 58 - Wasserrecht (animal protocols GZ: 342445/2016/12 and MA 58-221180-2021-16).

### Zebrafish husbandry

Zebrafish (*Danio rerio*) were raised according to standard protocols (28°C water temperature, 14/10 hour light/dark cycle). TLAB fish were generated by crossing AB and natural variant TL (Tupfel Longfin) zebrafish and used as wild type for all experiments. *MZoep* ^*-/-*26^ and *toddler* ^*-/-*10^ double mutant, as well as *tg(drl:gfp)*^28^, *toddler* ^*-/-*^ lines and *tg(gsc:CAAX-GFP)*^42^ wild-type lines were generated by crossing the two respective lines.

### Genotyping of mutants

Genotyping of *toddler* ^*-/-*10^ and *MZoep* ^*-/-*26^ mutants was performed by PCR amplification. As previously published ^10^, the *toddler* PCR amplicon (toddler_gt_F and toddler_gt_R) was digested with RsaI as the mutation destroys the corresponding restriction site. Detection of digest product (wild type: 192 + 318 nt, *toddler* ^-/-^: 510 nt) was performed using standard gel electrophoresis with 4% agarose gels. The *oep* PCR amplicon (oep_gt_F and oep_gt_R) was digested with Tsp45l as the mutation introduces the respective restriction site. Detection of the digested product (wild type: 300 nt, *oep* ^*-/-*^: 150 + 150 nt) was performed using standard gel electrophoresis with 4% agarose gels.

### Injection of mRNAs and morpholinos into zebrafish embryos

Capped mRNAs for *toddler, oep, aplnrb, aplnrb-sfgfp, cxcr4b, cxcl12a, lifeact-gfp, human h2b-bfp, human h2b-rfp* and *f’bfp* were transcribed from linearized plasmids using the SP6, T7 or T3 mMessage machine kit (Ambion), according to manufacturer’s protocol. All mRNAs were injected into 1-cell stage embryos unless indicated otherwise. All plasmids were previously described (Table S1).

Double knockdown of *aplnra* and *aplnrb*, as well as knockdown of *lefty2* were performed using morpholino (MO) injection as previously described^10,43–46^. Briefly, 0.5 and 1 ng of MOs against *aplnra* (cggtgtattccggcgttggctccat; GeneTools) and *aplnrb* (cagagaagttgtttgtcatgtgctc; GeneTools) respectively or 12.5 ng of MO against *lefty2* (agctggatgaacagagccatgac; GeneTools) were injected into 1-cell stage embryos. A control MO (cctcttacctcagttacaatttata; GeneTools) was used at equivalent concentrations.

**Table 1.** List of plasmids.

| Plasmid | mRNA | Restriction/Polymerase | Source |
| --- | --- | --- | --- |
| AP242 | toddler | BgIII/SP6 | Pauli et al., 2014 |
| R013 | oep | NotI/T7 | Zhang et al., 1998 |
| AP552 | aplnrb | EcoRV/T7 | Pauli et al., 2014 |
| AP606 | aplnrb-sfgfp | BgIII/SP6 | Pauli et al., 2014 |
| R160 | lifeact-gfp | XbaI/T3 | Raz Lab |
| R203 | human h2b-bfp | EcoRV/SP6 | Tsai et al., 2020 |
| R009 | human h2b-rfp | NotI/SP6 | https://www.addgene.org/53745/ |
| R202 | f'bfp | EcoRV/SP6 | Shindo et al., 2018 |

**Table 2.**
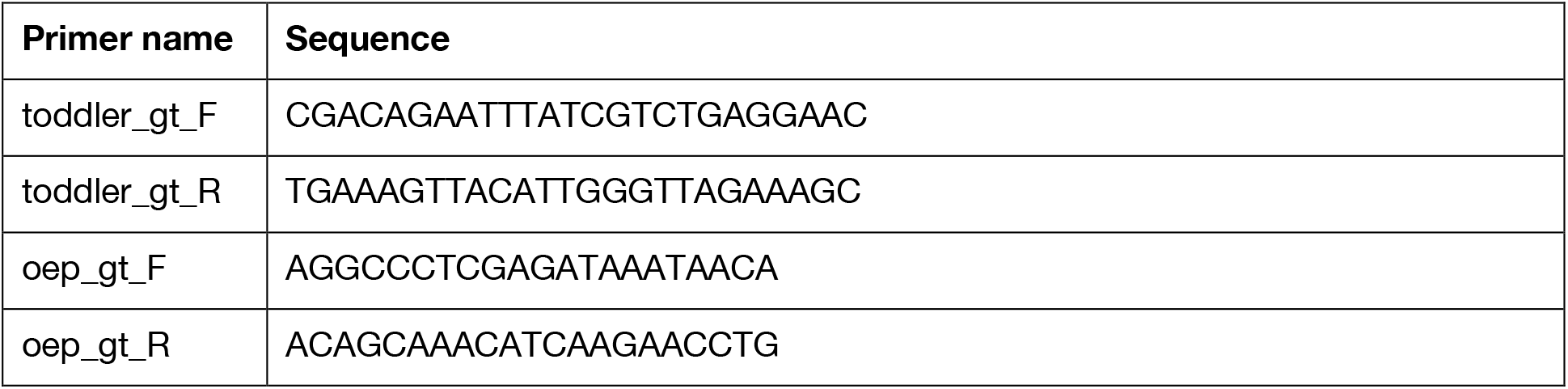
List of primers.

### Live cell imaging

#### Light sheet microscopy

For light sheet microscopy, four embryos were mounted in 0.6% low-melt agarose in 1XPBS in glass capillaries (Brand, 20 μL). The capillary was placed in the heated (27°C), with fish water filled sample chamber of a Zeiss Z1 light sheet microscope. The two most suitably oriented embryos were imaged.

To assess morphology and migration behavior of wild-type and *toddler* ^*-/-*^ cells, four successfully transplanted embryos (only one genotype per experiment) were mounted. The two embryos that had the best orientation (animal pole straight up or down + lateral orientation + transplanted cells in the center of the frame at the margin) were selected for imaging. Time laps series of Z-stacks were taken from both embryos over 5 hours (from sphere stage to end of gastrulation) with a 20x objective.

For global cell tracking experiments, wild-type *tg(drl:gfp)* or *toddler* ^*-/-*^ *tg(drl:gfp)* embryos were injected with 100 pg *h2b-rfp* mRNA at the 1-cell stage and mounted at sphere stage. The 0.6% low-melt agarose was supplemented with fluorescent beads (1:1000, TetraSpeck™ microspheres, 0.2 μm, fluorescent blue/green/orange/dark red, Invtirogen #T7280) that were used to adjust the light sheet offset and stabilize the imaging sequence. Four embryos were mounted, and the two most ideally oriented embryos (animal pole up or down) were selected for imaging. Time laps series of Z-stacks were taken from both embryos over 8 hours with a 10x objective. At 24 hours post fertilization, z-stacks from four positions around the embryo were acquired (0°, 90°, 180° and 270° in reference to the position of the time course imaging) to determine the location of dorsal and ventral sides within the embryo. Finally, 6 z-stacks of 1000 z-slices were acquired with lasers turned off (used to remove background during subsequent data analysis). Raw light sheet data was converted to tiff files. For cell morphology analyses, the data was binned 4x in xy to allow for better data handling. The region of interest (internalizing migrating cell) was identified in binned images and automatically cropped (in x/y/z and time) in the raw data, using a custom-made app (https://drive.google.com/drive/folders/1183kxnarTEyb1-4Otzyh0RlQhEnHO8Oo?usp=sharing). Cropped imaging files were analyzed as described below.

**Table 3.**
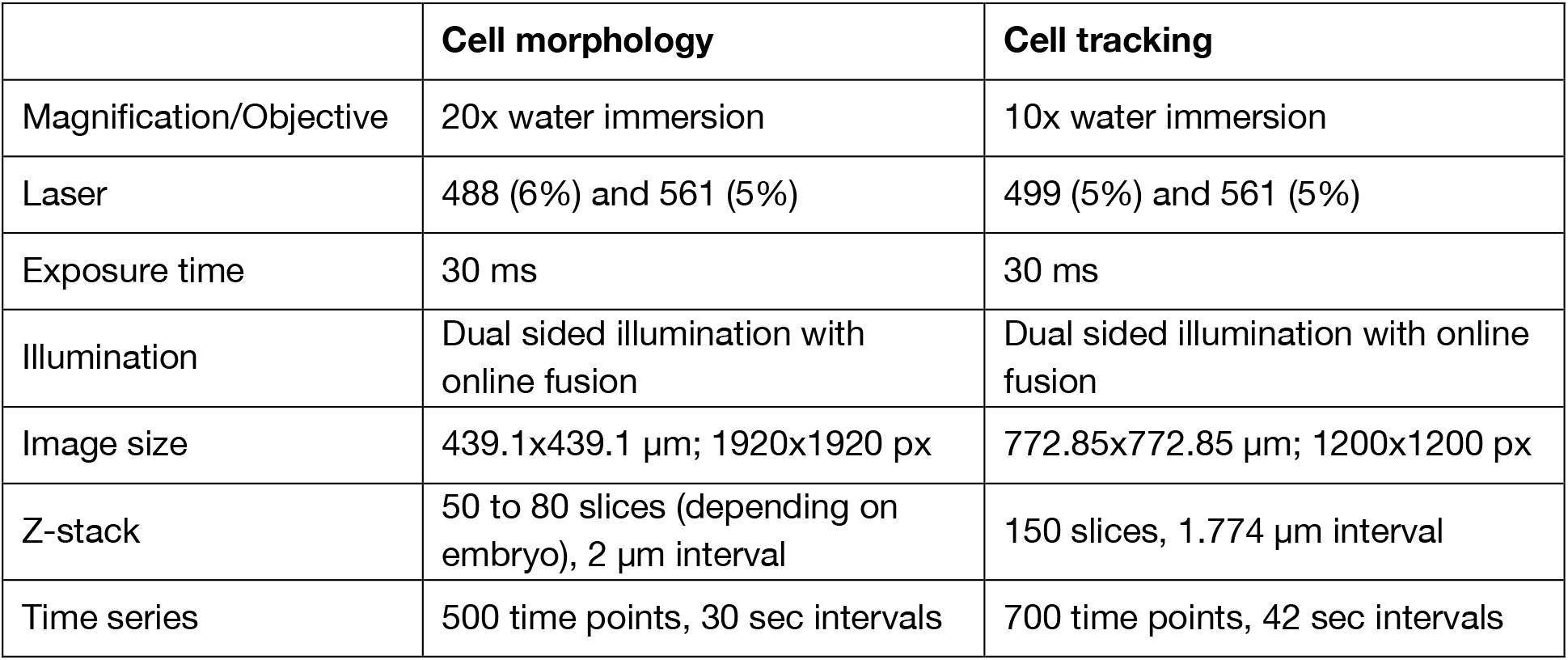
Settings for light sheet microscopy.

|  | **Cell morphology** | **Cell tracking** |
| --- | --- | --- |
| Magnification/Objective | 20x water immersion | 10x water immersion |
| Laser | 488 (6%) and 561 (5%) | 499 (5%) and 561 (5%) |
| Exposure time | 30 ms | 30 ms |
| Illumination | Dual sided illumination with online fusion | Dual sided illumination with online fusion |
| Image size | 439.1x439.1 $\mu\text{m}$; 1920x1920 px | 772.85x772.85 $\mu\text{m}$; 1200x1200 px |
| Z-stack | 50 to 80 slices (depending on embryo), 2 $\mu\text{m}$ interval | 150 slices, 1.774 $\mu\text{m}$ interval |
| Time series | 500 time points, 30 sec intervals | 700 time points, 42 sec intervals |

**Table 4.**
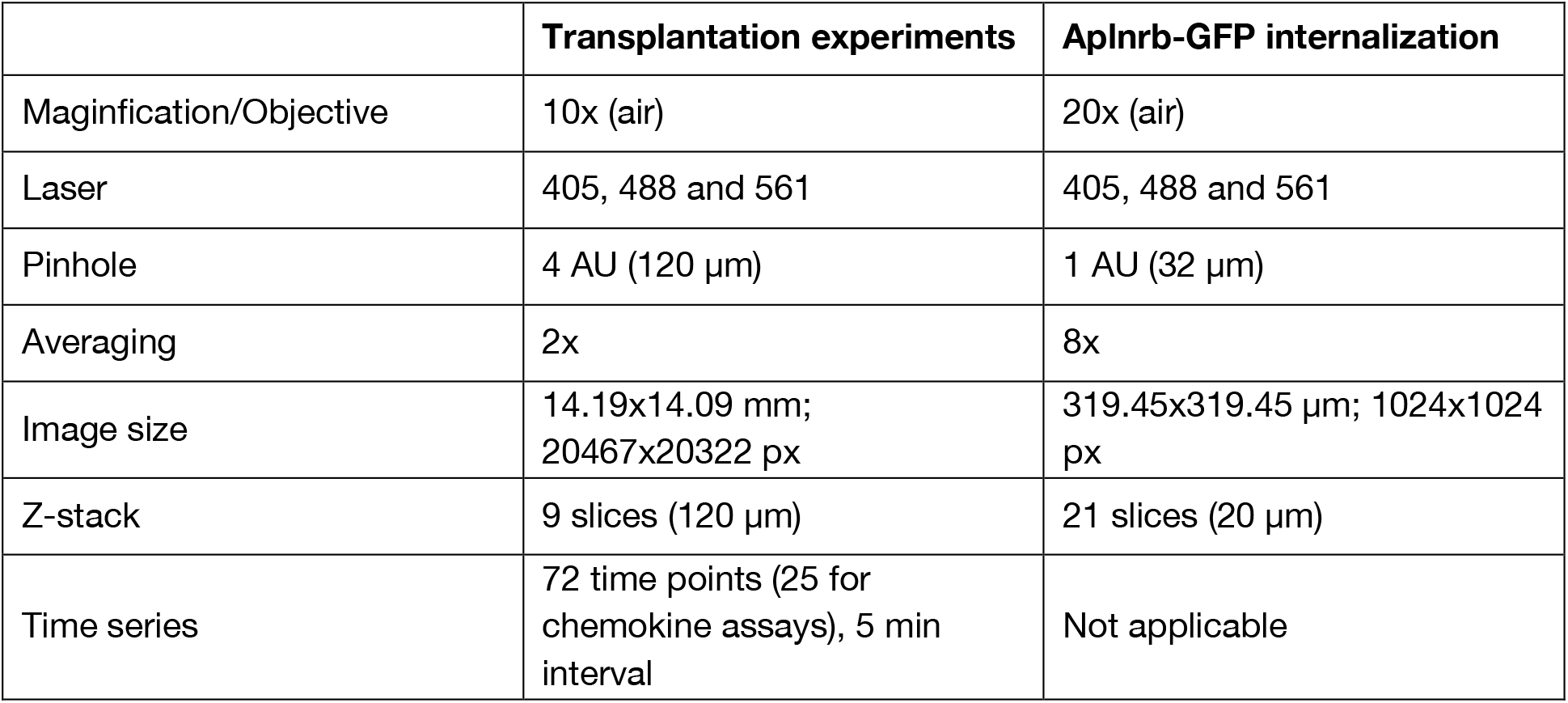
Settings for confocal microscopy.

|  | **Transplantation experiments** | **Aplnr-GFP internalization** |
| --- | --- | --- |
| Magnification/Objective | 10x (air) | 20x (air) |
| Laser | 405, 488 and 561 | 405, 488 and 561 |
| Pinhole | 4 AU (120 $\mu\text{m}$) | 1 AU (32 $\mu\text{m}$) |
| Averaging | 2x | 8x |
| Image size | 14.19x14.09 mm;<br>20467x20322 px | 319.45x319.45 $\mu\text{m}$; 1024x1024 px |
| Z-stack | 9 slices (120 $\mu\text{m}$) | 21 slices (20 $\mu\text{m}$) |
| Time series | 72 time points (25 for chemokine assays), 5 min interval | Not applicable |

#### Confocal microscopy

For confocal imaging, embryos were mounted in the desired orientation in a drop of 0.8% low-melt agarose in 1XPBS on round glass bottom dishes (ibidi). Agarose drops were left to harden before dishes were filled with E3 medium (5 mM NaCl, 0.17 mM KCl, 0.33 mM CaCl_2_, 0.33 mM MgSO_4_, 10^™ 5^% Methylene Blue). Time laps movies of z-stacks were acquired on an inverted LSM800 Axio Observer (Zeiss) with temperature incubation (27°C) for 6 hours. To assess Aplnrb-GFP internalization, z-stacks for each embryo were taken for only one time point at sphere stage.

### Transplantation assays

#### Cellular phenotype

To assess cell migration behavior of individual cells after internalization, donor embryos were injected with 100 pg *lifeact-gfp* mRNA. For light sheet imaging experiments, host embryos were injected with 100 pg *h2b-rfp* mRNA. Between 1 and 5 cells were taken from the marginal region of donor embryos at sphere stage and transplanted to margin of stage- and genotype-matched host embryos. At dome stage, embryos were mounted laterally with transplanted cells facing the glass dish and imaged on a confocal microscope or mounted in a glass capillary for light sheet microscopy and imaged as described above.

#### Cell autonomy

To assess the cell autonomy of Toddler signaling, embryos were prepared, and transplantations performed as described above. Donor and host embryos were stage-matched but of opposite genotypes (transplantation of wild type into *toddler* ^*-/-*^ and vice versa). Imaging and subsequent analysis were performed blindly.

#### Chemokine assay

All embryos used for the chemokine assay were *toddler* ^*-/-*^. Donor^receptor^ embryos were injected at the 1-cell stage with 150 pg *lifeact-rfp* mRNA only (control) or in combination with 200 pg *aplnrb-sfGFP* mRNA. Donor^ligand^ embryos were injected at the 1-cell stage with 100 pg Dextran-AlexaFluor488 only (control) or in combination with 100 pg *toddler* mRNA. Host embryos were injected at the 1-cell stage with 150 pg *h2b-bfp* mRNA. Embryos were left to develop until sphere stage. 50-100 cells were taken from donor^ligand^ embryos at and transplanted to the animal pole of stage-matched host embryos. 1-10 cells were taken from donor^receptor^ embryos and transplanted to the host embryo at positions surrounding the animal pole. Embryos were mounted on the animal pole and imaged as described above.

#### Localized Toddler source

Animal pole expression of Toddler was achieved by transplantation assays. Donor embryos were injected at the 1-cell stage with 100 pg Dextrane-AlexaFluor488 only (controls) or in combination with 200 pg *toddler* mRNA. Dextrane-AlexaFluor488 was used to trace successful injection and transplantation. At sphere stage, 50 to 100 cells were taken from donor embryos and transplanted to animal pole of stage-matched *toddler* ^*-/-*^ host embryos. For uniform expression of Toddler, 100 pg Dextran-AlexaFluor488 only (controls) or in combination with 2 pg of *toddler* mRNA were injected into 1-cell stage *toddler* ^*-/-*^ embryos. To achieve marginal expression, 100 pg Dextrane-AlexaFluor488 only (controls) or in combination with 10 pg *toddler* mRNA were injected into the yolk syncytial layer of 1k-cell stage embryos. Embryos were collected for in situ hybridization (see below) at 75% epiboly.

#### Sink removal

To test for sink function of mesendodermal cells and scavenger function of Aplnr, wild-type donor embryos were injected with 100 pg *lifeact-gfp* mRNA at the 1-cell stage. Host embryos were either untreated (*MZoep* ^*-/-*^) or injected with *aplnr* MOs to inhibit *aplnr* mRNA translation in wild-type embryos. 1 to 5 donor cells were taken from the margin at sphere stage and transplanted to the margin of a stage-matched host embryo. Transplantations of more than 10 cells were excluded from analysis to avoid the possibility of sink re-introduction. Embryos were mounted laterally with transplanted cells facing the objective and imaged using confocal microscopy as described above.

#### Toddler gradient

To assess the sufficiency of a Toddler gradient to guide cell migration in the absence of mesendodermal cells, transplantations were performed using *MZoep* ^*-/-*^, *toddler* ^*-/-*^ double mutant host embryos. Wild-type donor^reporter^ embryos were injected with 100 pg *lifeact-GFP* mRNA at the 1-cell stage. *MZoep* ^*-/-*^, *toddler* ^*-/-*^ double mutant donor^source^ embryos were injected at the 1-cell stage with 200 pg Dextran-AlexaFluor568 only (controls) or in combination with 200 pg *toddler* mRNA. 50 to 100 cells of donor^source^ cells were transplanted to the animal pole of a sphere stage host embryo. Subsequently, 1 to 5 LifeAct-GFP positive donor^reporter^ cells were transplanted to the margin of the same host. Embryos were mounted laterally with transplanted cells facing the objective and imaged using confocal microscopy as described above.

#### Single vs. collective cell migration

To generate a sink of Aplnr-expressing cells by transplantation, donor^reporter^ wild-type embryos were injected with 100 pg *lifeact-gfp* mRNA at the 1-cell stage. For donor^sink^, *tg(gsc:caax-gfp)* embryos were injected with 100 pg Dextran-AlexaFluor568 at the 1-cell stage. These embryos were used since they allow to identify and exclude the dorsal region at transplantation. Host and donor^sink^ embryos were injected with 0.5 ng of *aplnra* MO and 1 ng of *aplnrb* MO. At late dome stage, 50 cells from the ventral margin of donor^sink^ embryos followed by 5 to 10 cells from the margin of donor^reporter^ embryos were taken into the same needle and transplanted to the margin of stage-matched host embryos. Embryos were mounted laterally with transplanted cells facing the objective and imaged as described above. Imaging and subsequent analysis were performed blindly.

### Imaging analysis

#### Protrusions and polarity

Cells of interest for analysis were identified in the time-laps movies based on several criteria:

1. Ventral or lateral position in the host embryo: Cells that were transplanted to the dorsal side (as indicated by visible convergence and extension movements as well as the formation of somites at the end of the time-laps) were excluded from the analysis.
2. Only internalized cells were analyzed. Internalization was identified by a cell’s position in the z-dimension (lower cell layer) and a change in direction that indicates the transition from vegetally directed epiblast to animally directed hypoblast cell movement.
3. Sufficiently long cell tracks: Analysis of a cell had to be possible for at least 15 (light sheet; ≙30min) or 12 (confocal; ≙1h) time frames. Every cell that divided, moved out of frame or overlapped with another transplanted cell within this time window was excluded from analysis.
4. Analysis was performed before ∼70% epiboly (before the onset of convergence and extension movements).

Protrusions were distinguished based on the LifeAct-GFP signal of polymerized actin and defined by the following characteristics:

1. Lamellipodia: actin-mesh that expands several μm in width; wide protrusion that extends beyond the cell body.
2. Filopodia: thin actin strings that extend from the cell body.
3. Bleb: actin-free (marked by cytoplasmic, dispersed LifeAct-GFP), round membrane extension with polymerized actin at its base.

Each cell was analyzed from the time of internalization until one of the 4 criteria above was no longer applicable. In some cases, a cell divided shortly after internalization, therefore, analysis was started 10 (light sheet) or 2 (confocal) time frames (≙10min) after division to exclude mitosis-induced cell rounding and loss of polarity. Length and angle (in reference to animal pole) of protrusions were measured in ImageJ. A straight line through the center of the protrusion was drawn from the base (border to cell body) to the tip of the protrusion. A cell was counted as polarized if it was elongated and actin polymerization and protrusion formation was restricted to one side only, with additional protrusions (filopodia only) allowed within a 45° angle to either side. Protrusions were measured and counted for every cell in each time frame. Protrusion rate (protrusion/min) was calculated by normalizing the sum of all protrusions for one cell to the length of the time course (for light sheet) or the number of time frames (for confocal). To quantify orientation of polarization and protrusions, the numbers of angles within every 30° were counted and normalized to the total number of the same protrusion/polarity within the same genotype. As the position of dorsal and ventral in these embryos was not specifically determined, the left and right side of the resulting 360° rose plot were combined to be collectively counted as dorsal/ventral orientation.

#### Cell tracks

Cell tracks were obtained from maximum intensity projections of the cell of interest using tracking tools in Imaris (Imaris x64 9.7.2). First, time laps movies were corrected for epiboly movements. In light sheet data sets and chemokine assays, H2B-RFP/BFP-labelled cell nuclei were automatically tracked (autoregressive motion). Movies were then corrected based on 3 – 5 successful tracked nuclei, in close proximity to the cell of interest. For other confocal data, the margin (identified in bright field images) was manually tracked and used as reference for correction. Cells of interest in all confocal data (**Fig. 1-2 and 4, fig. S2-3 and S5-6**) and light sheet data in **Fig. 1** and **fig. S1** were manually tracked. Final tracks were exported and assembled using Adobe Illustrator. Average speed was calculated by dividing track length by track duration. Track straightness was calculated by dividing displacement by track length. Net displacement to the source was determined by measuring the vertical distance between start and end point of the track. The end point of the track was either the point at which the cell first made contact with the source, or, if no contact was made, after 2h of imaging.

#### Global cell tracking

Global cell tracking of all nuclei in the acquired light sheet microscopy data (**Fig. 3 and fig. S4**) was based on the approach of Jaqaman et al., 2018. To this end, detected nuclei were greedily interconnected into short tracklets, which were then merged to form the final tracks. The interconnection of both the detected nuclei and tracklets was computed by solving a linear assignment problem using LAPMOD, a sparse variant of the Jonker & Volgenant linear assignment solver^51^. Very short tracklets of less than 3 time points were pruned. New tracklets were not allowed to form within a sphere with a radius of 10 μm. Maximum allowed displacement between consecutive time points was set to 10 μm. Gaps between tracklets no longer than 3 time points were closed if the maximum displacement per time point stayed below a threshold of 10 micron.

The nuclei were detected using a multi-scale Laplacian of Gaussian filter (5 scales between 1 and 2.5 micron) after preprocessing the imaging data using the following operations: (1) A per-pixel mean image was computed and used for background subtraction to simplify cell tracking; noise and outliers were removed below and above the 50th and 99.99th percentile of intensity. (2) Working in the log domain, the volumes were downsampled 2x along x- and y-coordinates to obtain close-to-isotropic spacing; a 7×7 px median filter was applied for all z-planes independently. (3) Furthermore, to simplify the tracking, the sequence was stabilized using a translation transformation that was estimated at each time point from the locations of the beads around the embryo using RANSAC. In particular, aligning was started at a reference point, and bead locations were always pre-aligned at the current time point using past transformation; the beads locations fed into RANSAC estimation were selected as mutual nearest neighbors after this pre-alignment.

The global cell tracking was implemented in Python (https://drive.google.com/drive/folders/12Q36RQGpzItsEP4bC3OKYiWuM53ZC7sv?usp=sharing) using OpenCV, scikit-image and scikit-learn (48, 49). The tracker was optimized to take advantage of a SLURM-based computation cluster, however, the implementation also allows running locally on a single computer, however, the runtime might be prohibitive.

For visualization and analysis of the trajectories (https://drive.google.com/drive/folders/1GRgfTBL03aOvJ6FsMxBItVcMB6h2_cZc?usp=sharing), a sphere was fitted to the extracted nuclei locations^54^, with latitude and longitude coordinates and the radial distance from the center of the embryo. The reference sphere was aligned with the animal and vegetal pole matching the north and south pole, respectively. Afterwards the animal-vegetal axis and the dorsoventral axis were interactively recovered. This brought all the experiments into a common reference frame and allowed for simpler interpretation.

Cell internalization was determined by calculating each cell’s frame-by-frame difference in radius at the margin. The minimum of the average changes indicated a peak of internalization movements, which occurred between 30% and 50% epiboly in all datasets. This was used to align the recordings in developmental timing.

To isolate the tracks of mesodermal cells, nuclei colocalizing with the Drl:GFP signal were identified. In relation to the background fluorescence, a cell was considered Drl:GFP marker positive with a green fluorescence level higher than 96% of all measurements (z-score 1.8) for at least 50 frames.

The animal-to-vegetal pole velocity was measured specifically for ventrolateral mesoderm cells. This was calculated by the change in distance relative to the margin over a moving window of 10 min (14 frames). The distance to margin is the difference in latitude between the position of the cell and the position of the margin at the corresponding time point.

### Compensation of Toddler overexpression

Overexpression of Toddler was achieved at different levels by injecting 0 pg (water control), 5 pg or 10 pg of *toddler* mRNA into 1-cell stage wild-type embryos. Half of the embryos of each batch were additionally injected with *lefty2* MO to enhance Nodal signaling and thus increase the number of mesodermal cells. Embryos were collected for in situ hybridization at 75% epiboly (see below).

### In situ hybridization

In situ hybridization was performed as previously described^55^. Briefly, wild-type and *toddler* ^-/-^ embryos were fixed at 75% epiboly in 3.7% FA overnight and hybridized with a DIG-labelled antisense probe against *aplnrb*^10^. After BCIP/NBT/Alkaline-phosphatase-staining, embryos were dehydrated in Methanol and imaged in BB/BA on a Stemi 305 steromicroscope (Zeiss).

To quantify the mesoderm migration phenotype, embryos were imaged laterally at 75% epiboly. The spread of *aplnrb-*positive cells was measured in ImageJ on the ventral side and normalized to the distance between margin and animal pole to account for differences in epiboly progression between individual embryos. Data was normalized to the average spread in wild-type embryos.

To assess the mesodermal cell density profile of embryos, the gray values for a 100-pixel wide rectangle from animal pole to margin were measured using ImageJ and inverted (0=white, 255=black). Values for all 100 pixels in one row were averaged. The lowest value measured along the animal-margin axis was subtracted as background, and measurements were normalized to the highest value (=1). All background-subtracted pixel values were plotted as position along the animal-margin axis (margin=0, animal pole=1).

### Internalization of Aplnr

To assess Aplnrb-GFP internalization, *toddler* ^*-/-*^ embryos were injected at the 1-cell stage with 100 pg f’bfp mRNA and 50 pg aplnrb-sfgfp mRNA. At the 32-cell stage, 100 pg Dextrane-AlexaFluor568 only (controls) or in combination with 100 pg *toddler* mRNA were injected into a single blastomere. Embryos were left to develop until sphere stage and then mounted on the animal pole and imaged as described above. To quantify Aplnrb-GFP internalization, cell outlines were manually traced in ImageJ based on the f’BFP signal. Mean fluorescence intensity of Aplnrb-GFP was measured for membrane and cytoplasm and multiplied with the membrane and cytoplasmic area, respectively. The ratio of total membrane and cytoplasmic signal was used as a measure of the internalization rate.

### Toddler half-life assessment

To measure the half-life time of Toddler protein, *toddler* ^*-/-*^ or *MZoep* ^*-/-*^, *toddler* ^*-/-*^ embryos were collected and left to develop until 30% epiboly. To test for an Aplnrb-dependent increase in the Toddler degradation rate, *toddler* ^*-/-*^ embryos were injected with 50 pg *aplnrb* mRNA at the 1-cell stage. At 30% epiboly, 1 ng of Toddler peptide (GL Biochem: DKHGTKHDFLNLRRKYRRHN**C**PKKR**C**LPLHSRVPFP, cysteine residues in bold were crosslinked according to Ho et al., 2015) was injected into the cell cap. 10 cell caps were collected every 30 min for 3 hours through manual removal of yolk and flash frozen in liquid nitrogen.

Western blotting was performed according to standard protocols. Briefly, embryo caps were supplemented with 10 μL of 2x Laemmli buffer and boiled for 5 min at 95°C. SDS-PAGE was performed using Any kD™ Mini-PROTEAN® TGX™ Precast Protein Gels (BioRad), loading 10 embryo caps per lane. Blotting was performed using a wet-blot system (BioRad). Protein detection was achieved using the following antibodies: anti-Toddler (rabbit, 1:500, provided by the Reversade Lab^11^) and anti-tubulin (mouse, 1:10000, Sigma-Aldrich T6074).

### Computational modeling

Here, we provide additional details for the modelling of self-generated Toddler gradients during zebrafish gastrulation.

#### Position of the problem

We write the conservation equation for the concentration of mesoderm cells *m*(*x,t*) and secreted Toddler *T*(*x,t*) as a function of time *t* and position *x* along the animal-vegetal axis (where *x* = 0 is the position of the margin, and where we restrict ourselves to a one-dimensional description thanks to the radial symmetry of the problem):

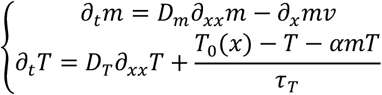

In this description, mesoderm cell concentration can change from i) free diffusion and ii) direction motion at speed *υ*, while Toddler concentration can change from i) free diffusion, ii) production (from ectoderm cells), iii) intrinsic degradation and iv) mesoderm consumption. We have denoted as *D*_*m*_ and *D*_*T*_ the diffusion coefficient of mesoderm cells (in the absence of directed motion) and Toddler molecules, *T*_0_(*x*) the target concentration of Toddler (which can be spatially modulated) in the absence of any mesoderm consuming it, *τ*_*T*_ the timescale of intrinsic Toddler degradation and *α* the consumption rate of Toddler from mesoderm cells (larger density of mesoderm cells signifying more receptor density for Toddler degradation—note that this implictly assumes that receptor density per cell is constant, an assumption that we relax at the end of the SI Note). To close this set of equation, we must additionally specify a dependency between directed cell migration velocity *υ* and Toddler concentration. How cells sense gradients is an area of active study, and different non-linear as well as adaptative responses have been uncovered in particular while interpreting GPCR signalling gradients (see for instance Jin 2013 for a review). Here, we explored two simple limits of gradient sensing: *υ* = *β∂*_*x*_*T* (i.e. cells move up an *absolute* gradient of Toddler) or 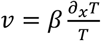 (i.e. cells move up a *relative* gradient in Toddler) - with *β* denoting in each case the strength of the coupling. We also note that this makes the important approximation (which we will come back to below) that Toddler gradients only impacts the average “advective” velocity of cells *υ* and not their random motility coefficient *D*_*m*_. This is a coarse-grained description which can arise from different microscopic models: for instance this doesn’t necessarily require the instantaneous speed of Toddler cells to be modified based on the Toddler, but could also be due to partial bias of their random walk based on the direction of cell polarity being biaised by the local Toddler gradient.

Finally, we specify boundary and initial conditions for this problem at the margin: Toddler and cells cannot escape out, leading to no-flux boundary conditions *∂*_*x*_*T*(*x* = 0) = 0 and *∂*_*x*_*m*(*x* = 0) = 0. For the initial conditions, although these are largely irrelevant for Toddler (given its constant renewal/production, the simplest is *T*(*x*, 0) = *T*_0_), they are key to specify for mesoderm cells. We first assume that the mesendoderm cell number is fixed and concentrated initially very close to the margin: *m*(*x*, 0) = *M*_0_*δ*(*x*, 0).

#### Key length and timescales in the problem

From these equations, two natural scales emerge: a time scale *τ*_*T*_ which represents the timescale of Toddler turnover, and a length scale 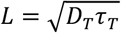 which represents the distance at which Toddler coming from a localized source is degraded. This is important for instance during our rescue experiments when we place Toddler-expressing cells at the animal pole: Toddler is predicted to decay exponentially from their location, at a length scale of *L*.

#### Parameter constraints, fitting and non-dimensionalization

A number of parameters can be constrained in this model. Toddler diffusion in particular can be calculated from the size of the Toddler molecule (around 4kDa) from the Stokes-Einstein relationship. For instance, Lefty, Cyclops or Squint, which have around 10 times the molecular weight of Toddler (and thus are expected to have 2-3 times the hydrodynamical radius) were shown to have free diffusion coefficients in zebrafish embryos of *D*_*T*_ ≈ 20*μm*^2^. *s*^−1^ (Muller et al, 2012). Therefore, we can estimate that *D*_*T*_ ≈ 50*μm*^2^. *s*^−1^ or *D*_*T*_ ≈ 3. 10^3^*μm*^2^. *min*^−1^. To estimate the time scale *τ*_*T*_ of Toddler degradation, we have made use of *MZoep* ^*-/-*^, *toddler* ^*-/-*^ double mutant embryos, which are devoid of both endogenous Toddler production and mesoderm-induced Toddler degradation. We injected 1 μg of Toddler peptide and performed a time course of Toddler degradation (**fig. S4**). This showed a roughly exponential decay, as predicted by our linear model, and from which we could extract *τ*_*T*_ ≈ 120 min (**fig. S4F-G**). Interestingly, this value is in the same order of magnitude as Lefty, Cyclops or Squint during early zebrafish embryo morphogenesis ^27^. Together, this predicts a length scale for Toddler propagation of 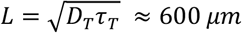, which is interesting as it is of comparable scale to the embryo itself (note that modelling the full complexity of diffusion in a cell-fluid mixture would give rise to slightly different pre-factors^58^, although this would have little quantitative impact on the predictions). This is consistent with the ability of margin cells to sense Toddler-expressing cells far away at the animal pole, to restore animal-directed migration in the experiments of **Fig. 2 and 4**.

Next, we examined the movements of cells transplanted in a *toddler* ^-/-^ background. As these cells show no measurable polarization/directed motion *υ* = 0, we reasoned we could use these experiments to constrain the value of free cell diffusion *D*_*m*_. We found that these cells diffused on a length scale of approximately 25*μm* from the margin during the 30 min of the timescale (**fig. S4K-L**), leading us to a rough estimate of *D*_*m*_ ≈ 20 *μm*^2^. *min*^−1^.

Once we rescale all time scales by *τ*_*T*_, all length scales by *L*:

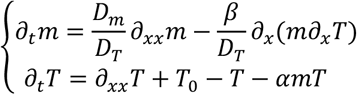

We can further rescale Toddler by this maximal concentration *T*_0_, and mesoderm by its initial total amount *M*_0_ leading to

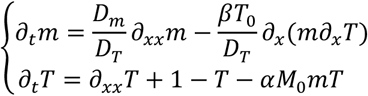

Thus, in addition to the length and time scales (which have been independently measured), this shows that the problem now only depends on 3 rescaled parameters: the relative diffusion coefficients of mesoderm and Toddler 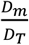 (which is strongly constrained by our measurements, and found to be very small), the rescaled consumption of Toddler from mesoderm *αM*_0_, and the rescaled coupling from

Toddler gradient to directed mesoderm speeds 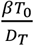. These two last parameters are harder to independently measure and should be therefore be considered as fitting parameters in the theory.

Importantly, however, analysis of the system of equations above finds that these last two parameters can largely be coarse-grained into a single one, with their product being the most relevant parameter: this is because *α* controls how strong of a gradient of Toddler is created, and *β* how strongly this gradient is interpreted, so that high *α* – low *β* and low *α* – high *β* give rise to similar velocities. Numerical simulations keeping the product *αβ* constant, but changing each by several orders of magnitude confirmed this (see **fig. S4C** where we multiply *β* by 5 and divide *α* by 5, giving nearly identical results to **fig. S4A**), although this effect breaks down at very high *α* (when mesoderm consumption of Toddler is so strong that the Toddler concentration reaches values close to zero). To check that we were not in this extreme regime of parameters, we performed additional experiments, repeating the Toddler degradation experiment, but in either *toddler* ^*-/-*^ (without *MZoep* ^*-/-*^, i.e. where Toddler is not produced, but mesoderm is present to degrade it) or *aplnrb*^*OE*^ (overexpression of Aplnr in the Toddler mutant background, i.e. no production of Toddler, and enhanced Aplnr-dependent degradation of Toddler) – see **fig. S4F,G** for details. In *toddler* ^*-/-*^ (compared to *MZoep* ^*-/-*^, *toddler* ^*-/-*^), we found very similar kinetics of Toddler degradation (*τ*_*T*_ ≈ 101min), which is consistent with only a small fraction of cells degrading Toddler within the emrbyo and constraining the value of 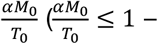 we take it equal to 1 throughout out simulations, although it is the product *αβ* which matters, see below). Even in the *aplnrb*^*OE*^ condition, we found only moderately faster Toddler degradation (*τ*_*T*_ ≈ 61min), confirming our hypothesis, and arguing that we can use the product *αβ* as the grouped single fitting parameter. Toddler concentration never reaching very small values also means that the two modes of gradient sensing that we discussed above (relative gradient 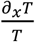 or absolute gradient *∂*_*x*_*T*) give similar predictions.

Another approximation of the model that we wished to verify was that the random motility of cells (represented by the diffusion coefficient *D*_*m*_ was unaffected by the local Toddler gradient. To verify this, we analyzed quantitatively the experiments presented in **Fig. 2**, in which cells with or without Aplnrb-sfGFP were transplanted at a distance from a Toddler-expressing or Toddler-deficient source. We thus quantified the average displacements of cells over 15min, either in the direction of the source (y-axis) or perpendicular to it (x-axis), and generated probability distributions for each case. As expected from a purely random and diffusive process in the x-direction, all three conditions displayed Gaussian distributions in step size centered around zero average displacement (**fig. S4M-N**). Importantly, the standard deviation (proportional to *D*_*m*_) was nearly identical in all three cases, arguing that random motility is unaffected by either the presence of a Toddler gradient or Aplnrb expression. Interestingly, when looking at the same distribution in the y-direction (towards the source), we found again that the standard deviation of the displacement was comparable between conditions (and also to its value in the x-direction, as expected from a random walk, **fig. S4M-N**). The only difference for the Toddler+Aplnrb condition was that the best-fit Gaussian distribution was not centered around 0, but instead around a non-zero average velocity value of 0.3 *μm*/*min* – as expected for a biased random walk and our model where Toddler gradients only act on the advective velocity *v* (**fig. S4M-N**).

Finally, although the model as defined above assumes that all cells internalize at the same time (initial Delta function at x=0 in the initial condition *m*(*x*, 0) = *M*_0_*δ*(*x*, 0)), the experimental situation is more gradual, with numbers of internalizing cells showing a broad temporal peak with typical variance *σ* of an hour ^13^. This can easily be taken into account by assuming that the initial condition is now zero mesoderm cells *m*(*x*, 0) = 0, but adding a gradual source term in the conservation equation for mesoderm cells: 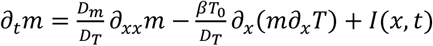, where *I*(*x, t*) is the spatio-temporal dynamics of internalization. Thus, we take in the simulations 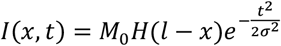: internalization only occurs close to the margin (represented by a Heaviside function decaying at *l* = 50*μm*), and on timescales *σ* = 1*h* ^13^. Although these are the parameters that we show in Fig. 4, we also ran simulations with the previous initial condition (synchronous internalization of all mesendoderm cells and found very similar results for both WT and Toddler mutant simulations, see **fig. S4A-E**).

Thus, in the following, we only fit *αβ* in the theory (simulations from main text are made for *αM*_0_ = 1). *αβ* is essentially proportional to the speed of migrating cells up a self-generated gradient. As we found an average speed of *υ* ≈ 0.08 *μm*/*min*, this means *αβ* ≈ 10^4^ in our unit simulations.

With the model fully parametrized in this manner, we turned to its predictions on a number of non-trivial features, such as the spatiotemporal density/velocity profiles of mesoderm migration in wild type and *toddler* ^*-/-*^ mutant, or transplantation assays (**Fig. 3, 6G, S7A,B**).

#### Model predictions

We first consider the case of the *toddler* ^*-/-*^ mutant, in which directed migration is negligible (*υ* = 0). Because the margin constitutes a hard boundary, internalized cells are expected to still migrate upwards to some degree according to a diffusive process with coefficient *D*_*m*_. Given our estimate of *D*_*m*_ from short-term trajectories, we thus asked how much cells were predicted to travel by pure diffusion (1D along the animal-vegetal axis) in the Δ*t*=3.5 hours between internalization and the 75% epiboly stage (which gave in the simulations around 200 μm – compared to 600 μm for wild type, **fig. S4K**) Interestingly, we measured the intensity profiles of mesoderm markers in *toddler* ^*-/-*^ compared to the wild type – as a proxy for mesoderm concentration along the animal-vegetal axis (**fig. S4H-J**), and found that they decayed around twice as fast in *toddler* ^*-/-*^ compared to wild-type embryos (we note that uncertainty in the exact diffusion coefficient of mesoderm cells, or the possibility of some small, non-zero directionality being retained in *toddler* ^*-/-*^ mutants embryos could explain the slightly stronger phenotype in the model).

In the presence of self-generated gradients, the system organizes into a travelling-wave solution, as expected from the literature ^23^, where cells adopt a non-zero net polarity/velocity towards the animal pole, as observed experimentally (**Fig. 3E and G-H**). Although this self-organized collective migration is robust to the details of the parameters, such as the effective diffusion length scale *L* for Toddler, such parameters do have an effect on the detailed spatiotemporal profiles of mesoderm migration. For very local Toddler diffusion (small *L*), only a few cells at the very edge sense the self-generated gradient, which cause them to initially migrate very fast compared to the front ones (see **fig. S4D** for a simulation with *τ*_*T*_ = 10*s*, so that the length scale *L* is around 20 μm, i.e. the cell size). However, this creates a concentration gradient of mesoderm cells, itself causing a concentration gradient of Toddler (which doesn’t come from diffusion, but rather from gradient of degradation from the gradient of mesoderm). Thus, cells can still migrate in a self-generated manner, although the cellular density gradient is more pronounced than the “front-like” solutions shown in **Fig. 3** (limit of large effective Toddler diffusion relevant here as the length scale *L* is of order of the embryo size as described above).

Comparing these predictions to our tracking data of mesoderm cells (marked by *drl:GFP*) undergoing post-internalization migration towards the animal pole, we found similar qualitative features, with largest velocities at the edge, which decreased both as a function of time and distance from the edge. More quantitatively, we compared kymograph for cellular velocity as a function of position as a distance from margin and time (**Fig. 3H**) – note that we show in the kymograph the total effective velocity of cells as measured by tracking, which is the sum of the advective velocity and the diffusive flux (which can have directional contribution in the presence of a density gradient). The total flux of cells reads as *J_tot_* = *mβ∂*_*x*_*T* − *D*_*m*_*∂*_*x*_*m* where the first term is the advective contribution proportional to local Toddler gradient and the second is the diffusive flux. We thus define *υ*_*tot*_ = *J*_*tot*_/*m* = *β∂*_*x*_*T* − *D*_*m*_*∂*_*x*_*m*/*m* as the total average velocity of mesoderm cells at position *x*, which is what is plotted in **Fig. 3H**. Importantly, although for the wild type, we predict that the contribution of diffusion is rather small compared to advection (see **fig. S4E** for a simulation with zero mesendoderm free diffusion, *D*_*m*_ = 0), the latter becomes dominant in *toddler* ^*-/-*^, where advection is close to zero.

#### Effect of number of transplanted cells on the resulting dynamics

As discussed in the main text, the mechanism of self-generated gradients that we propose relies on a collective effect, where the number of cells at the margin matter, as they dictate the strength of the gradient-shaping. To explore further the effect on sink function and cell number, we refined the model to include different cell types: regular Aplnr-expressing mesoderm cells *m*(*x, t*), which degrade Toddler, and cells which do not degrade Toddler *m*_0_(*x, t*). The equation on *m*(*x,t*) is exactly the same as before, as is the equation on Toddler (only *m*(*x,t*) participate in Toddler degradation, while *m*_0_(*x,t*) does not enter the Toddler equation. The equation on *m*_0_(*x,t*) reads 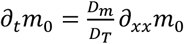 (i.e. no directed migration term). As we show in **Fig. 4H**, simulating transplants of small numbers of wild-type cells in *aplnr*^*MO*^ embryos (large *m*_0_ density, low *m* density) resulted in little migration, while transplants of small numbers of wild-type cells in wild-type density was effectively the same as regular wild-type migration (as transplanted cells are identical to the surroundings). On the other hand, simulating large clusters of wild-type cells in *aplnr*^*MO*^ embryos (large *m*_0_ density, intermediate *m* density) resulted in an intermediary phenotype (**Fig. 4H**), as seen in the data (**Fig. 4**).

#### Overexpression of Toddler and Apelin receptor

We next consider the effect of overexpression of Toddler, which has shown to give rise to defects in upward migration ^10^. This is modelled by a change in the baseline production of Toddler *T*_0_:

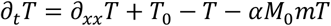

Where previously we had non-dimensionalize the problem to *T*_0_ = 1. When considering different *T*_0_, here, the assumption of absolute vs. relative gradient sensing (resp. *v* = *β∂_x_T* or 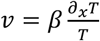) do impact on the resulting prediction: for absolute gradient sensing, the gradient (and thus migration) increases for increasing *T*_0_, whereas it is almost insensitive to it for relative gradient sensing. However, it should be noted that the equation above implicitly assumes that mesoderm cells (via their Apelin receptors) can take up arbitrary amounts of Toddler ligand. Although this may be correct for the wild type, this situation might not generically hold for overexpression phenotypes, especially if Apelin receptors are internalized with Toddler. To take this later feature of GPCR signalling into account, we supplemented the equations above with a conservation equation for the concentration of Apelin receptors within a cell *r*(*x, t*):

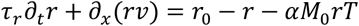

Note that there are no diffusion terms in this equation, as Apelin receptors don’t spatially exchange between cells, although they are “transported” spatially together with the movements of mesoderm cells (advective term putting *r* in the co-moving frame of cells). This first-order equation assumes that there is a baseline equilibrium of Apelin receptor at the membrane (time scale of recycling *τ*_r_, equilibrium concentration *r*_0_), but that each event of Toddler internalization also removes an Apelin receptor, which is the same sink term as in the Toddler equation, now rewrote to depend also on *r*:

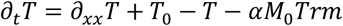

Note that the later term also is multiplied by the mesendoderm concentration *m*, as we have defined *r*(*x, t*) as the single-cell concentration of Apelin receptors.

When simulating these equations with *r*_0_ = 1 and *τ*_2_ = 30min (although these values had little bearing on the results), we found very similar dynamics to previously, with Apelin receptor concentration very slightly increased at the back of the front (where Toddler concentration is lower). However, when simulating a 10x increase in Toddler production (*T*_0_ = 10, which was close concentrations previously reported to induce a Toddler overexpression phenotype ^10^ **(fig. S7C)**, we found that this now resulted in an impaired migration (**fig. S7A**), which was due to a much lower concentration of *r* (due to Toddler saturation), which was less able to make Toddler gradient. The more the Toddler production was increased, the stronger the defect in migration was. However, increasing the production of Apelin receptor *r*_0_ could restore the normal wild-type phenotype (**fig. S7B**). The exact value necessary to exactly compensate depended on the turnover rate of Apelin receptors, for *τ*_r_ = 30min, *r*_0_ = 2.25 was required to restore the normal migration profile from *T*_0_ = 10. Interestingly, this closely matches previous experimental observations of an epistatic relationship between Toddler and Apelin receptors (**fig. S7C**).

**Fig. S1.**
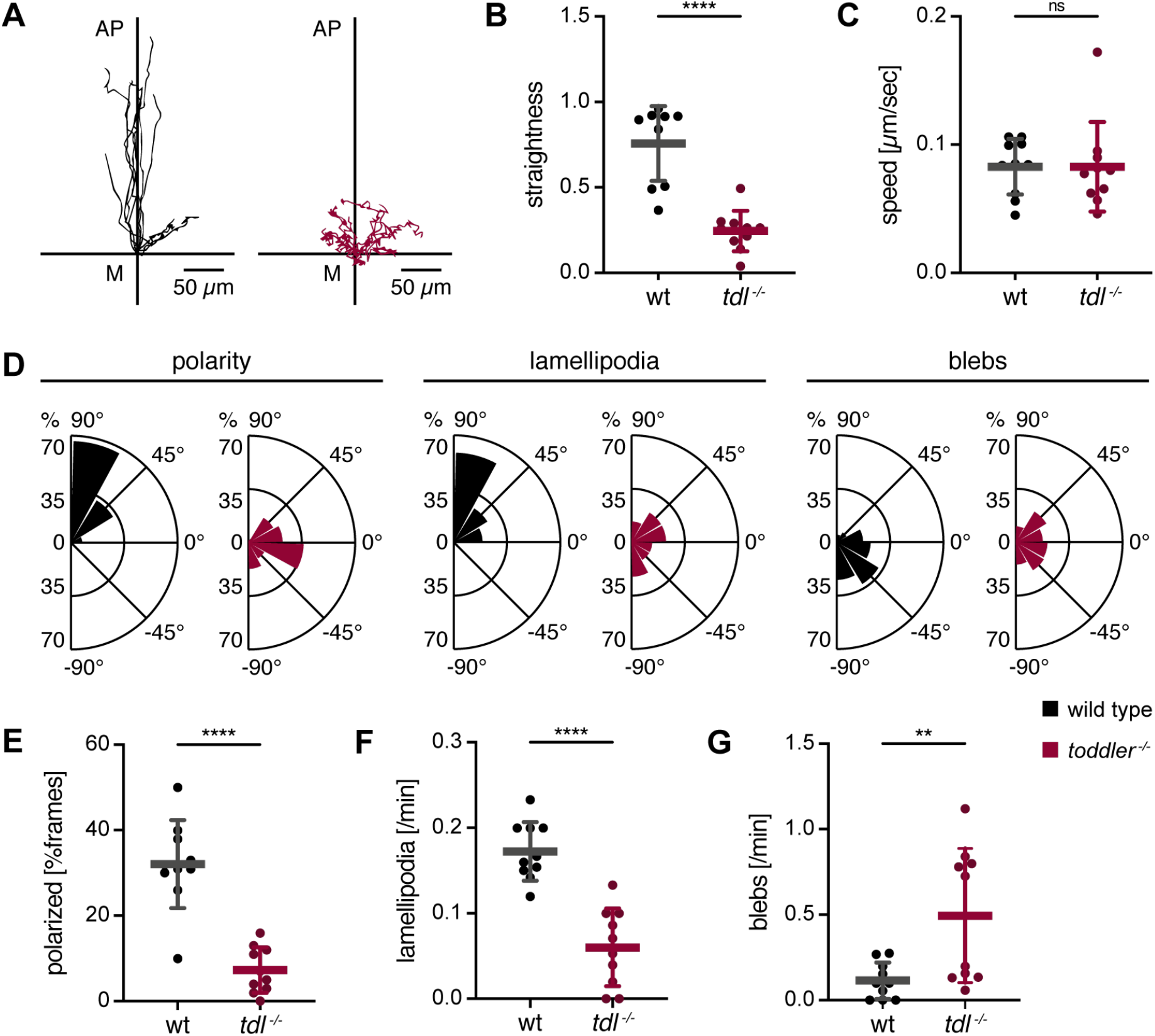
Mesendodermal progenitors are unpolarized and lack animal pole-directed protrusions in the absence of Toddler. Cell transplantation assays to assess the migration behavior of mesendodermal progenitors in the presence versus absence of Toddler signaling using light sheet microscopy. LifeAct-GFP-labelled reporter cells were transplanted from the margin of a wild-type or *toddler* ^*-/-*^ donor embryo to the margin of a stage-and genotype-matched host embryo. **(A)** Tracks of wild-type (left) and *toddler* ^*-/-*^ (right) reporter cells. Cells were tracked for 30 min after internalization. x-axis = margin; y-axis = animal-vegetal axis; coordinate origin = start of track. **(B)** Quantification of track straightness. **(C)** Quantification of migration speed. **(D)** Rose plots showing relative enrichments (percentages) of orientations of polarity, lamellipodia and blebs normalized to the total number of polarity axes or respective protrusions of all cells within the same genotype. **(E)** Quantification of cell polarity represented as the percentage of frames in which a cell was polarized. **(F)** Quantification of lamellipodia. Data represents newly formed lamellipodia per minute. **(G)** Quantification of blebs. Data represents newly formed blebs per minute. Data are means ± standard deviation (SD). Significance was determined using unpaired t test; ****, p < 0.0001; **, p < 0.01; n.s., not significant. n = 10 cells. Wild type (black); *toddler* ^*-/-*^ (red). Rose plots: 90° = animal pole; 0° = ventral/dorsal; −90° = vegetal pole. All images and graphs are oriented with the animal pole towards the top.

**Fig. S2.**
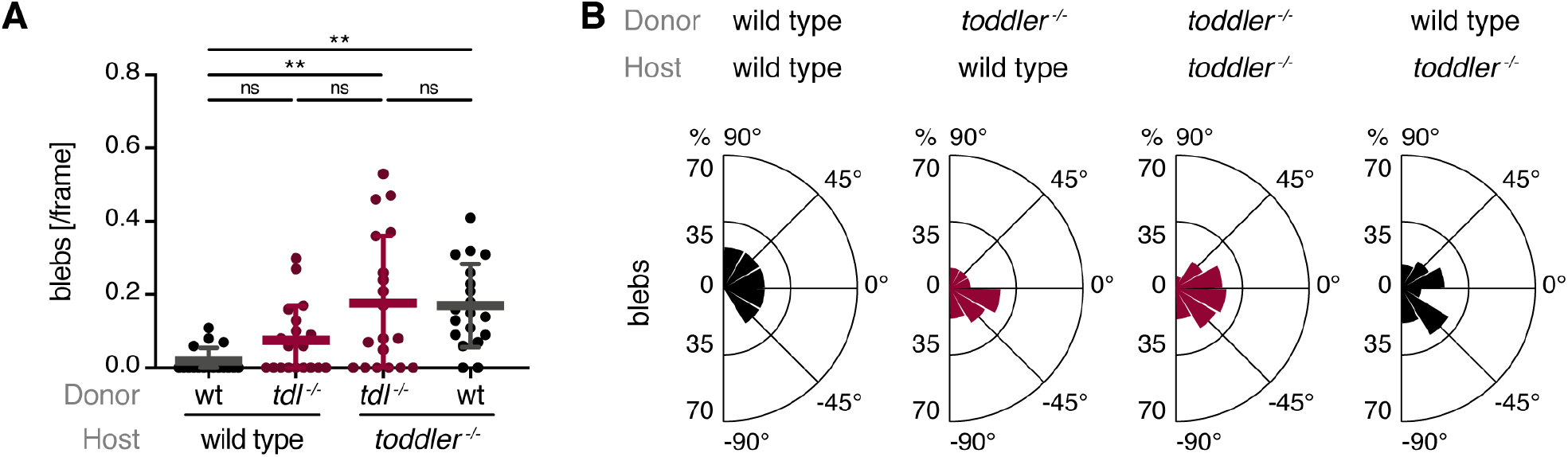
*toddler* ^*-/-*^ cells display an increase in cell blebbing. Cell transplantation assays to assess the cell autonomous or non-autonomous regulation of cell blebbing by Toddler signaling. **(A)** Quantification of blebs represented as the average number of blebs detected per frame (see Materials and Methods for classification of blebs). **(B)** Rose plots showing relative enrichments (percentages) of orientations of blebs, normalized to the total number of blebs of all cells within the same condition. Rose plots: 90° = animal pole; 0° = ventral/dorsal; −90° = vegetal pole. Data are means ± SD. Significance was determined using unpaired t test; **, p < 0.01; n.s., not significant.

**Fig. S3.**
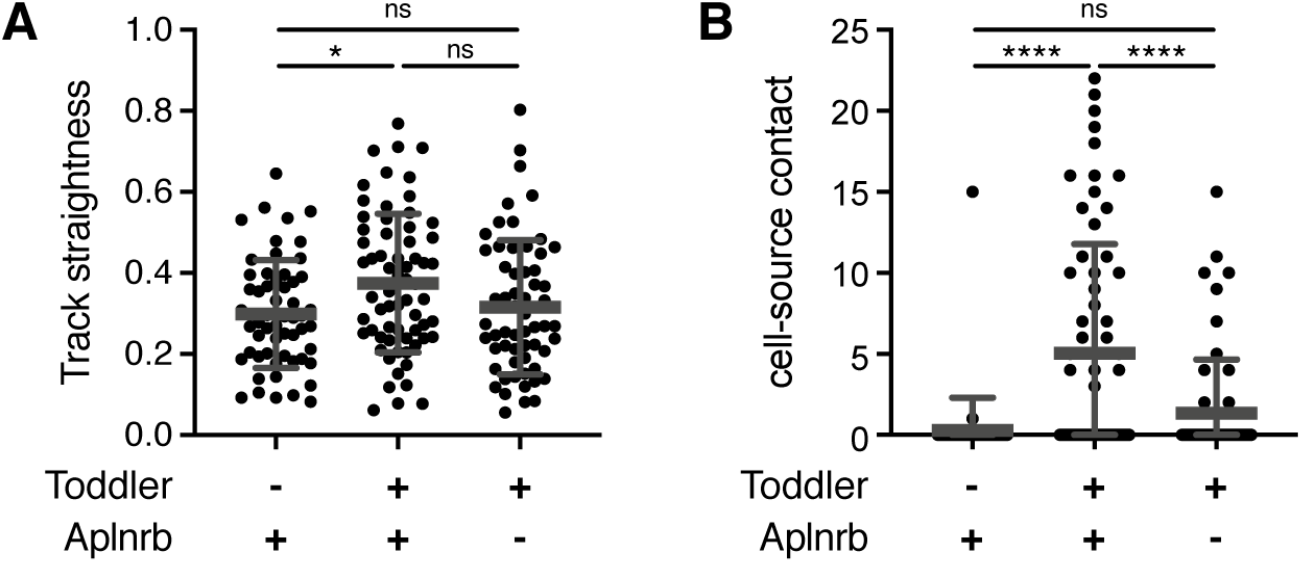
A localized source of Toddler attracts Aplnrb-sfGFP-expressing cells. Assessment of Toddler’s ability to act as a chemoattractant for Aplnr-expressing mesodermal cells. **(A)** Track straightness of all cells imaged in Fig. 2B. Track straightness was calculated based on full 120 min tracks as displacement divided by track length, irrespective of when the cells encountered the source. **(B)** Quantification of cell-source contact. The longest consecutive streak of frames in which contact with a Toddler source cell was detected, was plotted for each cell. n = 56, 65 and 59 respectively. Data are means ± SD. Significance was determined using one-way ANOVA with multiple comparison; ****, p < 0.0001; *, p < 0.05; n.s., not significant.

**Fig. S4.**
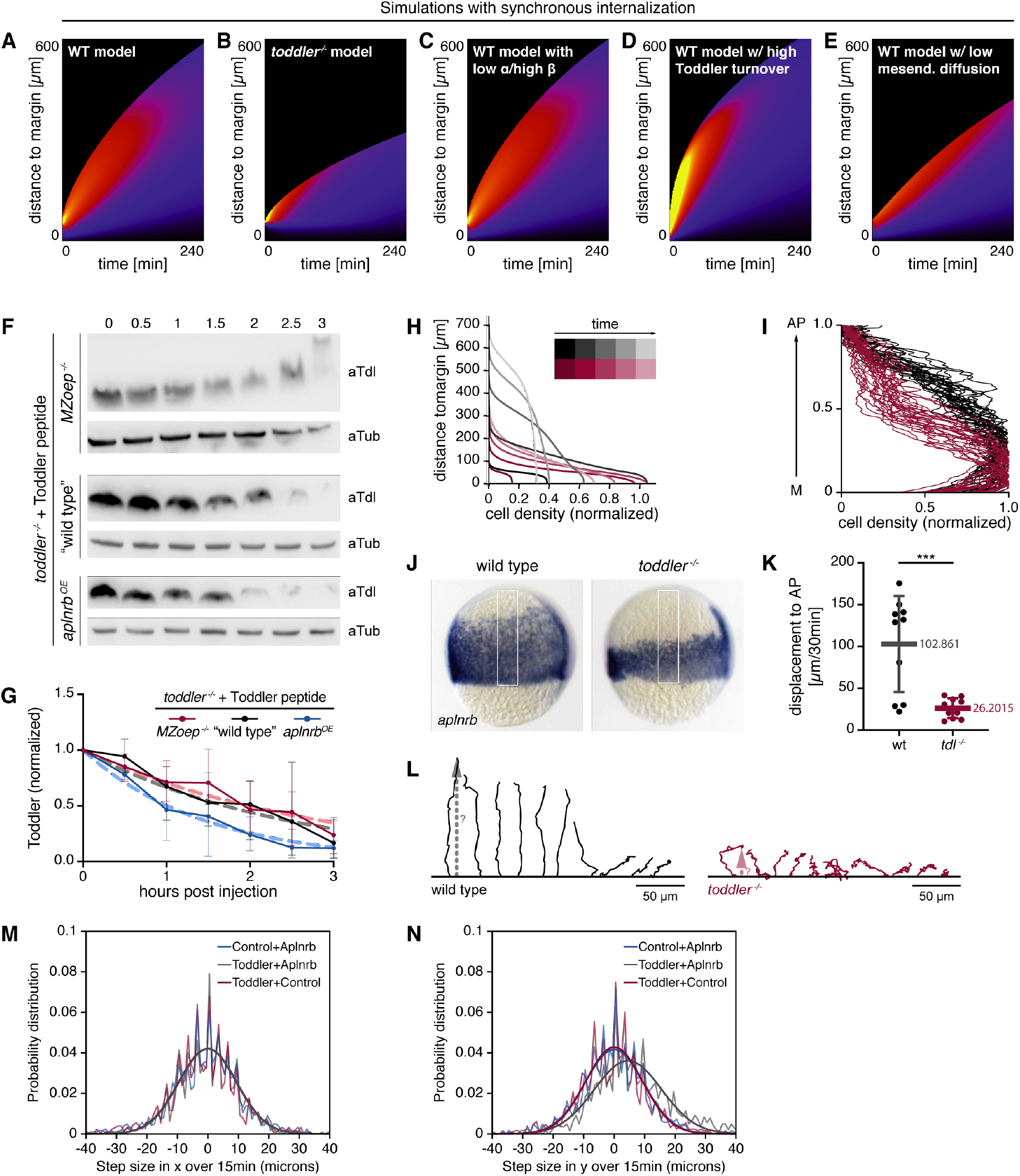
Determination of parameters for computational modeling. **(A-E)** Kymographs for normalized cellular velocities as a function of time (x-axis) and distance from the margin (y-axis): Sensitivity analysis of the model to different assumptions shown in the case of synchronous internalization from the onset (see Materials & Methods for details). **(A-B)** Simulations with the same simulation parameters as Fig. 4H for WT (**A**) and *toddler*^*-/-*^ (**B**) embryos but for synchronous internalization, showing qualitatively similar dynamics. **(C)** Simulations keeping the product *αβ* constant but multiplying *β* (coupling between local Toddler gradients and velocity) by 5 and dividing *α* (sink strength) by 5 gives rise to similar dynamics compared to panel (A). **(D)** Simulations with the same model parameters as (**A**) but with faster Toddler baseline degradation *τ*_*T*_ = 10*s*, decreasing the range of Toddler gradient propagation. **(E)** Simulations with the same model parameters as (**A**) but with negligible mesendoderm random cell motility *D*_*m*_ = 0, giving rise to a smaller range of migration but qualitatively similar velocity profiles. **(F-G)** Assessing Toddler peptide stability in the presence and absence of Aplnr. **(F)** Western Blot analysis of the Toddler peptide degradation rate. *In vitro* synthesized Toddler peptide was injected into *MZoep* ^*-/-*^ (top, deficient of mesodermal cells), *toddler*^*-/-*^ (middle, wild-type levels of Aplnr expression), and *aplnr*^*OE*^ (bottom, overexpressing Aplnr) embryos. Embryos were collected every 30 min for 3 hours and used for Western Blot analysis, probing for Toddler (aTdl) and alpha-Tubulin (aTub; loading control). **(G)** Quantification of Toddler levels (normalized to Tubulin) at different time points. Dotted lines represent exponential fits for degradation curves that were used to calculate Toddler half-life in the different conditions. **(H-I)** Measurements of cell displacement towards the animal pole to determine Toddler-independent cell velocity in *toddler* ^*-/-*^ embryos. **(H)** Predicted spatiotemporal profile of mesoderm cell densities in the wild-type (grey) and *toddler*^*-/-*^ (red) condition. **(I-J)** Experimental assessment of mesodermal cell density along the animal-margin axis. **(I)** Quantification of cell density along animal-margin axis in wild-type (black) and *toddler* ^*-/-*^ (red) embryos. n = 25 embryos. **(J)** Images for *in situ* hybridization for *aplnrb* of a representative wild-type (left) and *toddler*^*-/-*^ (right) embryo. White box indicates area measured for quantification **(K)** Quantification of the net animal pole (AP)-directed displacement based on tracks in **(I)**. Data are means ± SD. Significance was determined using unpaired t test; ***, p < 0.001. **(L)** Individual migration tracks of cells presented in fig. S1. The perpendicular net displacement of each cell from the margin (dotted line) was measured using the end point of each cell track 30 min after internalization. Track start is at the margin. Animal pole is shown towards the top. Wild type (black), *toddler* ^*-/-*^ (red). **(M-N)** Distribution of step size, defined as distance migrated by cells in the x (M) and y (N) directions within 15 minutes (x-direction: left-right movement in respect to the cell-source axis; y-direction: movement towards the source; see **Fig. 2A-C** for schematics and plotting of the trajectories), across all three conditions examined (Aplnrb-sfGFP-expressing cells migrating towards a Toddler-expressing cell, Aplnrb-sfGFP-expressing cells migrating towards a Toddler-negative control cell, Aplnrb-deficient control cells migrating towards a Toddler expressing source). All datasets are well-fitted by Gaussian distribution 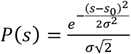 (black lines), as predicted from a biased random walk, with the only non-zero bias *s*_0_ occurring in the y-direction for the Aplnrb+Toddler condition. Best-fit variance: *σ* = 9.50, 9,49, 9.47*μm* for resp. blue, grey and red datasets in panel J. Best-fit variance: *σ* = 9.32, 11.05, 9.57*μm* for resp. blue, grey and red datasets in panel K, with a best-fit bias *s*_0_ = 4.55*μm* for the Toddler+Aplnrb condition.

**Fig. S5.**
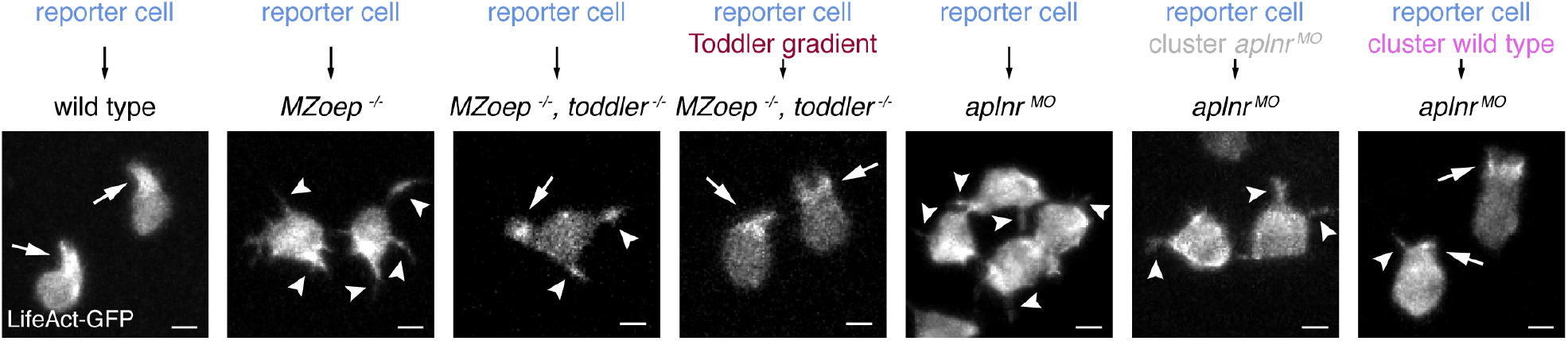
Mesodermal cells form ectopic protrusions in the absence of a Toddler sink. Representative confocal images of time-lapse series of transplanted reporter cells in the presence or absence of a mesodermal sink or Toddler gradient, as depicted in **Fig. 4A**. Arrows and arrowheads indicate lamellipodia and filopodia, respectively.

**Fig. S6.**
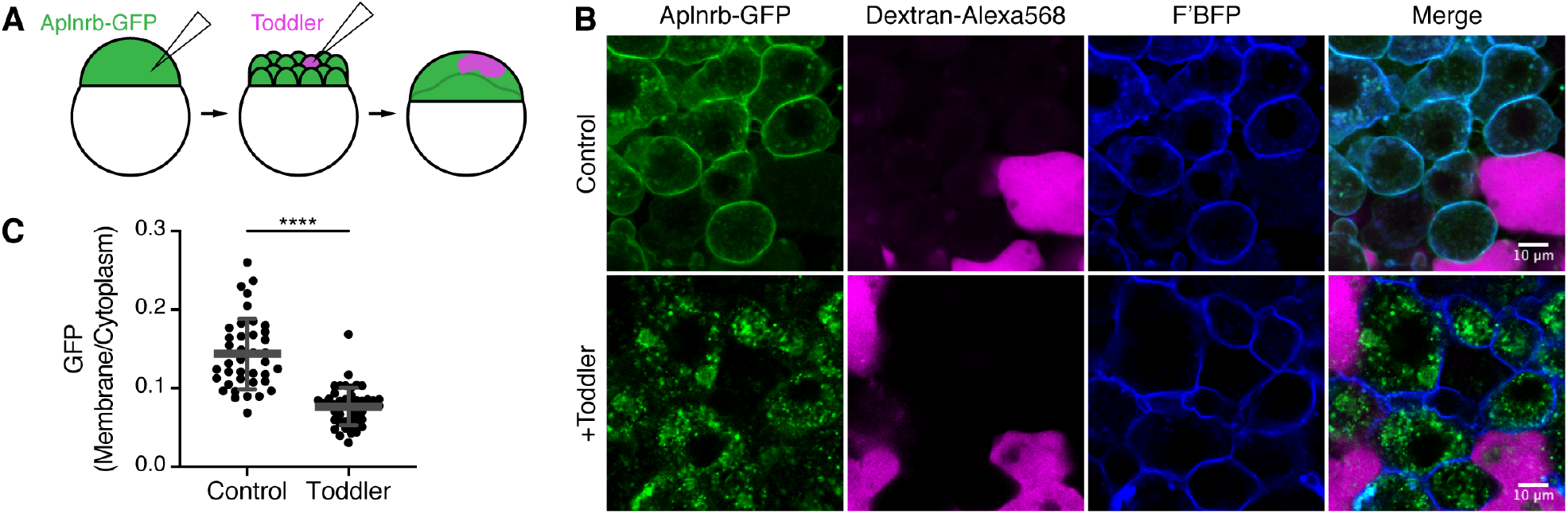
Aplnr internalizes upon Toddler signaling. **(A)** Schematic representation of experimental set-up. *Aplnrb-GFP* mRNA (green) was injected together with *F’BFP* mRNA (farnesylated, membrane-tethered blue-fluorescent protein) into 1-cell stage embryos to be ubiquitously expressed. Dextran-AlexaFluor568 alone or together with *toddler* mRNA was injected into one blastomere of a 32-cell stage embryo to achieve mosaic expression. **(B)** Representative confocal images of Aplnrb-GFP localization in the absence (top, n = 32 cells) or presence (bottom, n = 27 cells) of Toddler. **(C)** Quantification of the subcellular localization of Aplnrb-GFP. Data are means ± SD. Significance was determined using unpaired t test; ****, p < 0.0001.

**Fig. S7.**
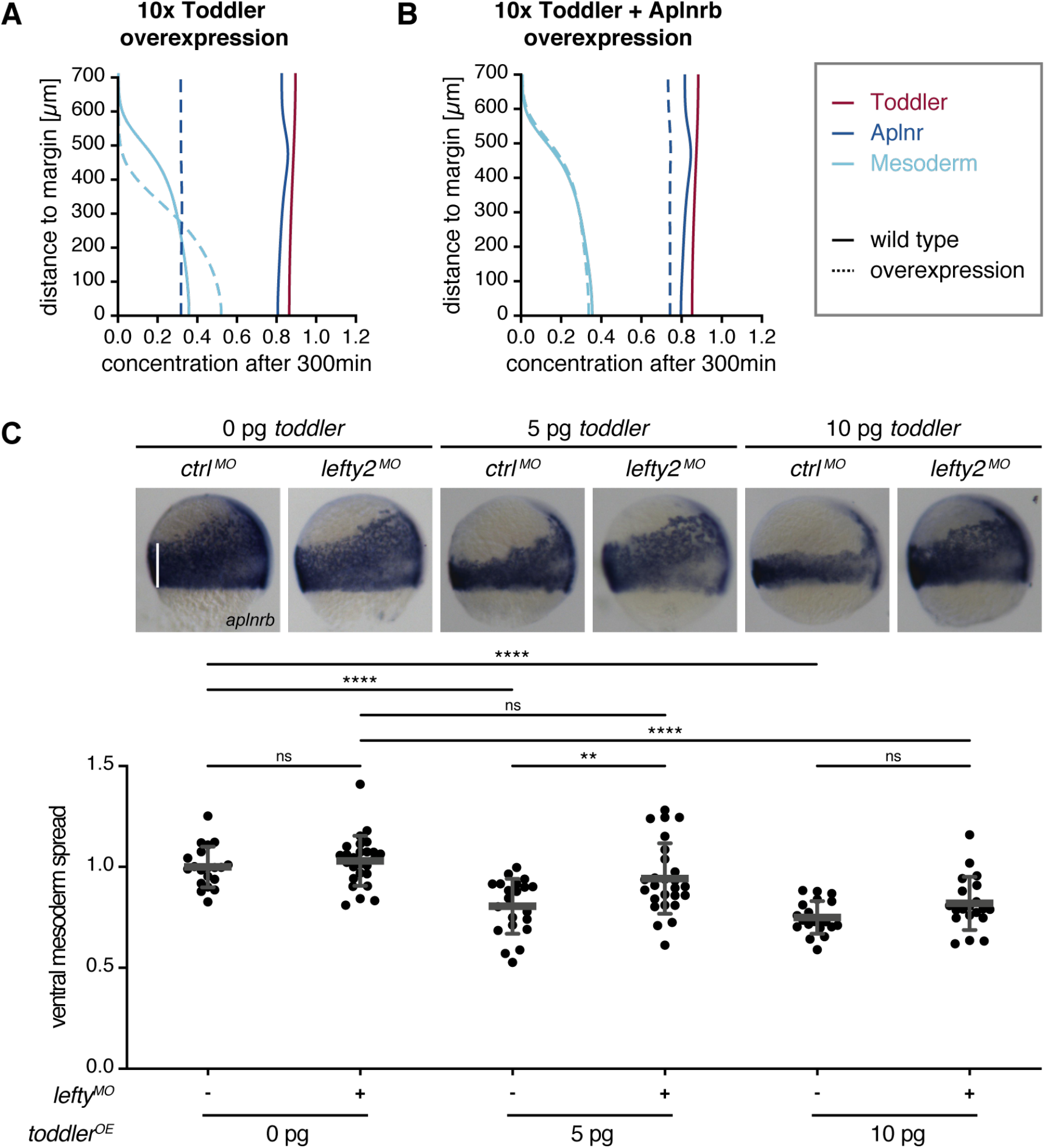
Overexpression of Toddler disrupts self-generated gradient but can be compensated by increasing the number of mesodermal cells. **(A)** Numerical simulation of mesoderm (light blue) migration upon Toddler (red) overexpression (10-fold, i.e. T_0_ = 10), assuming a finite capacity of Aplnr (dark blue) to remove Toddler and cells sensing relative Toddler gradients (see **Materials and Methods** for details). Toddler over-expression causes a decrease in Aplnr concentration, which prevents efficient Toddler gradient formation, resulting in slower mesoderm migration. Unbroken lines represent wild-type scenario, dotted lines depict the changes upon Toddler overexpression. **(B)** Numerical simulation of mesoderm migration upon Toddler and Aplnr overexpression (same parameters as in (A) for Toddler; Aplnr overexpression by 2.25-fold, see **Materials and Methods** for details), which rescues normal mesoderm migration. Representation as described in (A). **(C)** Experimental confirmation of the simulation presented in (A-B). Toddler was overexpressed at different concentrations in wild-type embryos (rescuing concentration of *toddler* mRNA injection into 1-cell stage *toddler*^*-/-*^ embryos is 2 pg) injected with control or *lefty2* MO to assess the compensation of increased Toddler levels upon increase of mesodermal cells (reducing the levels of the Nodal inhibitor Lefty2 increases the amount of Aplnr-expressing mesodermal cells). (Top) Representative *in situ* hybridization images using *aplnrb* as a probe to detect mesodermal cells. Embryos are shown as lateral views, with dorsal on the right. The white vertical line was used to measure the mesoderm spread from the margin towards the animal pole. (Bottom) Quantification of the mesoderm spread in each condition relative to the average mesoderm spread in wild-type embryos. Data are means ± SD. Significance was determined using one-way ANOVA with multiple comparison; ****, p < 0.0001; **, p < 0.01; n.s., not significant.

## Supplementary Movie Legends

**Movie S1** | **Internalized wild-type cells polarize and extend actin-rich lamellipodia towards the animal pole**. Light sheet time-lapse imaging (time interval: 1 min) of LifeAct-GFP-labelled wild-type cells transplanted to the margin of a wild-type host embryo. Establishment of a polymerized actin network and lamellipodia is marked by accumulation of LifeAct-GFP at the front of the cell. Movie starts after cells have successfully internalized and shows efficient animal-pole directed migration of these cells. Each frame is a maximum intensity projection of a z-stack. Animal pole is to the top. Scale bar, 10 μm.

**Movie S2** | **Internalized *toddler*** ^***-/-***^ **cells fail to polarize and fail to form actin-rich lamellipodia**. Light sheet time-lapse imaging (time interval: 1 min) of LifeAct-GFP-labelled *toddler* ^*-/-*^ cells transplanted to the margin of a *toddler* ^*-/-*^ host embryo. Cells display a loss of polarization and lamellipodia formation, as well as the formation of ectopic filopodia around the cell periphery. Movie starts after cells have successfully internalized. Cells fail to move towards the animal pole. Each frame is a maximum intensity projection of a z-stack. Animal pole is to the top. Scale bar, 10 μm.

**Movie S3** | **A subset of internalized *toddler*** ^***-/-***^ **cells displays an increased blebbing phenotype**. Light sheet time-lapse imaging (time interval: 1 min) of LifeAct-GFP-labelled *toddler* ^*-/-*^ cells transplanted to the margin of a *toddler* ^*-/-*^ host embryo, revealing the lack of actin-rich protrusion. Instead, actin-deficient blebs are formed as observed in a subset of analyzed *toddler* ^*-/-*^ cells. Movie starts after cells have successfully internalized. Cells fail to move towards the animal pole. Each frame is a maximum intensity projection of a z-stack. Animal pole is to the top. Scale bar, 10 μm.

**Movie S4** | **Confocal imaging of internalized wild-type cells**. Confocal time-lapse imaging (time interval: 5 min) of LifeAct-GFP-labelled wild-type cells transplanted to the margin of a wild-type host embryo, confirming morphology and animal pole-directed migration observed in movie S1. Each frame is a maximum intensity projection of a z-stack. Animal pole is to the top. Scale bar, 50 μm.

**Movie S5** | **Confocal imaging of internalized *toddler*** ^***-/-***^ **cells**. Confocal time-lapse imaging (time interval: 5 min intervals) of LifeAct-GFP-labelled *toddler* ^*-/-*^ cells transplanted to the margin of a *toddler* ^*-/-*^ host embryo, confirming morphology and lack of animal pole-directed migration observed in movie S2. Each frame is a maximum projection of a z-stack. Animal pole is to the top. Scale bar, 50 μm.

**Movie S6** | ***toddler*** ^***-/-***^ **cells display normal morphology and migration to the animal pole when transplanted into a wild-type host embryo**. Confocal time-lapse imaging (time interval: 5 min) of LifeAct-GFP-labelled *toddler* ^*-/-*^ cells transplanted to the margin of a wild-type host embryo. Each frame is a maximum projection of a z-stack. Animal pole is located towards the top. Scale bar, 50 μm.

**Movie S7** | **Wild-type cells display defects in protrusion formation, polarization and migration to the animal pole when transplanted into a *toddler*** ^***-/-***^ **host embryo**. Confocal time-lapse imaging (time interval: 5 min interval) of LifeAct-GFP-labelled wild-type cells transplanted to the margin of a *toddler* ^*-/-*^ host embryo. Each frame is a maximum projection of a z-stack. Animal pole is located towards the top. Scale bar, 50 μm.

**Movie S8** | **Aplnr-expressing cells are attracted by a localized source of Toddler**. Confocal time-lapse imaging (time interval: 5 min) of Aplnrb-sfGFP-expressing cells reacting to an ectopic Toddler source. Left: Aplnrb-expressing cells (blue) are placed next to a control source (grey). Middle: Aplnr-expressing cells (blue) are placed next to a Toddler-overexpressing source (red). Right: Aplnr-deficient cells (grey) are placed next to a Toddler-overexpressing source (red). Mesodermal and source cells are labelled with LifeAct-GFP and Dextran-AlexaFluore568, respectively. Each frame is a maximum projection of a z-stack. Source is located towards the top. Scale bar, 20 μm.

**Movie S9** | **Wild-type reporter cells lose directional migration and polarity in *MZoep*** ^***-/-***^ **host embryos**. Confocal time-lapse imaging (time interval: 5 min interval) of wild-type reporter cells (blue) transplanted into *MZoep* ^*-/-*^ embryos to test for the necessity of a mesendodermal Toddler sink. Left: Reporter cells transplanted to the margin of an *MZoep* ^*-/-*^ host embryo, which is deficient of mesendodermal progenitor cells. Middle: Reporter cells transplanted to the margin of an *MZoep* ^*-/-*^, *toddler* ^*-/-*^ double mutant host embryo, which is deficient of mesendodermal progenitor cells and Toddler expression. A control source (grey) was transplanted to the animal pole. Right: Reporter cells transplanted to the margin of an *MZoep*^-/-^, *toddler* ^*-/-*^ double mutant host embryo, which is deficient of mesendodermal progenitor cells and Toddler expression. A Toddler source (red) was transplanted to animal pole to mimic an ectopic Toddler gradient. Each frame is a maximum projection of a z-stack. Animal pole is located towards the top. Scale bar, 50 μm.

**Movie S10** | **Wild-type reporter cells lose directional migration and polarity in *MZoep*** ^***-/-***^ **host embryos**. Confocal time-lapse imaging (time interval: 5 min interval) of wild-type reporter cells (blue) transplanted into *aplnr*^*MO*^ embryos to test for the effect of individual versus collective cell migration in an environment of ubiquitous Toddler levels. Left: Reporter cells transplanted to the margin of an *aplnr*^*MO*^ host embryo, which forms mesoderm but is deficient in Aplnr expression. Middle: Co-transplantation of reporter cells and a control cluster of Aplnrb-deficient cells (grey) to the margin of an *aplnr*^*MO*^ host embryo. Right: Co-transplantation of reporter cells and a cluster of Aplnrb-expressing cells (magenta) to the margin of an *aplnr*^*MO*^ host embryo to mimic sink activity. Each frame is a maximum projection of a z-stack. Animal pole is located towards the top. Scale bar, 50 μm.

